# CalciumInsights: An Open-Source, Tissue-Agnostic Graphical Interface for High-Quality Analysis of Calcium Signals

**DOI:** 10.1101/2025.06.04.657923

**Authors:** Deiver Suarez-Gomez, Norma C Perez-Rosas, Gabriel I Miranda-Contreras, Santiago R Colom-Braña, Weiwei Zhang, Mayesha Sahir Mim, Shelly Tan, David Gazzo, Adrian Buganza Tepole, Qing Deng, Gregory T Reeves, Clara E Isaza-Brando, Christopher J. Staiger, David Umulis, Jeremiah Zartman, Mauricio Cabrera-Ríos

## Abstract

Fluctuations and propagation of cytosolic calcium levels at both the cellular and tissue levels show complex patterns, referred to as calcium signatures, that regulate growth, organ development, damage responses, and survival. The quantitative analysis of calcium signatures at the cellular level is essential for identifying unique patterns that coordinate biological processes. However, a versatile framework applicable to multiple tissue types, allowing researchers to compare, measure, and validate diverse responses and recognize conserved patterns across model organisms, is missing. Here, we present a post-processing tool, CalciumInsights, which leverages the R packages Shiny and Golem. This tool has a graphical user interface and does not require software programming experience to perform calcium signal analysis. The open-source software has a modular framework with standardized functionalities that can be tailored for various research approaches. CalciumInsights provides descriptive statistical analysis through various metrics extracted from dynamic calcium transients and oscillations, such as peak amplitude, area under the curve, frequency, among others. The tool was evaluated with fluorescence imaging data from three model organisms: *Danio rerio*, *Arabidopsis thaliana*, and *Drosophila melanogaster*, demonstrating its ability to analyze diverse biological responses and models. Finally, the open-source nature of CalciumInsights enables community-driven improvements and developments for enabling new applications.

**Author Summary:** This manuscript introduces CalciumInsights, an open-source tool for calcium signature analysis. Designed to be a versatile tool that works with various tissue types and biological systems, CalciumInsights has an easy-to-use graphical user interface. Our program simplifies metrics extraction while maintaining the quality of the analysis by integrating several algorithms. CalciumInsights stands out for its user-friendliness, ease of use, and robust data exploration features, such as tunable filters for improved accuracy. These features promote inclusivity and lower barriers to scientific research by making calcium signature analysis accessible to users of all programming skill levels.

## Introduction

Cytosolic calcium is a crucial second messenger that facilitates communication within and between eukaryotic cells, coordinating activities across different cellular compartments (1,2). For example, these activities include but are not limited to the following biological processes: exocytosis, contraction, metabolism, transcription, fertilization, growth and development of multicellular organisms, immunity, and wound healing (3–6).

As a universal second messenger in almost all biological contexts, understanding the commonality of oscillation patterns or wave propagation across divergent systems is an essential step toward recognizing and defining the Rules of Life, which are fundamental principles governing biological processes and behaviors (4,7–10). Cellular and tissue dynamics of cytosolic Ca^2+^ are complex, encompassing intracellular waves, spikes, oscillations, and coordinated cell-to-cell transmission (2,11,12). Diverse fluorescent, genetic, and imaging technologies are employed to acquire time-series data of Ca^2+^ fluctuations within cells and tissues. These tools include aequorin (13), fluorescent Ca^2+^ indicator dyes (14), genetically encoded calcium reporters or indicators (15,16), fluorescence and confocal microscopy (17), two-photon microscopy, electrophysiological techniques, optogenetic tools (18), and high-speed imaging systems.

By utilizing dedicated software and hardware, precise and reliable time-lapse series that accurately depict the dynamics of Ca^2+^ within target cells or tissues in multiple organisms can be generated. Determining the region of interest (ROI) where the Ca^2+^ signals occur is essential for a thorough quantitative analysis. Both manual and automated segmentations are frequently employed. Typically, the former is done with image analysis software such as Fiji (19). On the other hand, automated ROI segmentation can analyze the system’s spatiotemporal properties and extract the ROI without human intervention, accelerating the process and reducing the potential human error (20,21). Following the segmentation of cells or identification of ROIs, fluctuations in fluorescence known as Ca^2+^ transients can be extracted for further analysis. Standard computational tools like MATLAB, Python, R, or Java (22–29) can be used to analyze Ca^2+^ transients; however, these programming platforms provide a barrier for many biological applications due to a steep learning curve, especially for researchers lacking the necessary computational background. As such, there is a compelling need for enhanced functionality in novel, readily available tools to extend the analytical potential of the previously outlined methods to a broader user base. There are proprietary software tools like CytoSolver (30), Clampfit (31), Igor Pro (32), CyteSeer (33), and HARVELE that facilitate the analysis of the Ca^2+^ transients (34–36). However, open-source techniques enhancing the development of tools that are as powerful but more widely accessible are still lacking (37). To address this deficiency, we developed CalciumInsights. This analysis tool is designed to be user-friendly, featuring an interactive graphical user interface (GUI) that eliminates the need for programming knowledge or the purchase of proprietary software with complex and unintuitive functions (38). CalciumInsights leverages the Golem R-package to create modular Shiny applications in R (39,40), enabling streamlined development and allowing users to incorporate custom-built modules. This integration fosters collaboration within the open-source community, as users with advanced computational skills can tailor the application more precisely to their specific model system. These tailored applications can then be shared and utilized by others, expanding the utility and adaptability of CalciumInsights across diverse research contexts.

In this work, we used time-lapse fluorescence microscopy images from three common model organisms --*Danio rerio* (zebrafish), *Arabidopsis thaliana* (plant), and *Drosophila melanogaster* (fruit fly) -- to assess the efficacy and broad applicability of CalciumInsights. The Ca^2+^ traces from these images were analyzed and characterized using the metrics described in material and methods section. This versatility highlights CalciumInsights as an adaptable tool for analyzing data from different organisms across various imaging modalities and spatiotemporal resolutions, without bias toward any specific biological system or response.

## Materials and Methods

### Mathematical background

Ca^2+^ signals are essential in many scientific and medical settings because they provide vital insight into cellular activity involved in physiological processes. However, noise and artifacts often hinder the precise interpretation of these signals, complicating accurate analysis (41). Therefore, extracting meaningful information from these signals is essential for a comprehensive understanding of Ca^2+^ activity. To address this, we employ the Loess approach, a variation of *Locally Weighted Regression* (42,43). This method adjusts local polynomials to the data with weighted modifications to facilitate signal noise modeling and attenuation (44–46). The outcome is a smoothed representation of the original analog signal. The metrics of interest are then deduced from this smoothed signal. It is important to note that Loess parameters must be carefully chosen based on the characteristics of the experimental data and the specific metrics of interest.

#### Definition of the Loess Method

The fundamental mathematical basis of Loess involves estimating a smoothed value 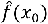 at a point *x*_0_ through a weighted combination of observed values *y*_*i*_in its vicinity (42,43). Mathematically, this is expressed as:

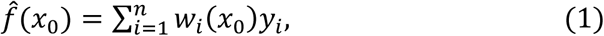

where *w*_*i*_(*x*_0_) are the weights assigned to each *y*_*i*_around *x*_0_, depending on the relative distance between *x*_0_and *x*_*i*_, which is controlled by the hyperparameter *span* ∈ [0,1]. The span determines the neighborhood of points considered in the fitting process by setting the size of the window used for the local regression. The width of the window is determined so that the proportion of data points it includes, relative to the entire dataset, matches a specified span parameter. A small span gives more weight to nearby points, allowing for a more flexible and responsive adjustment to local variations in the Ca^2+^ signal. In contrast, a large span assigns a more uniform weight to a greater number of points, resulting in a smoother adjustment less sensitive to local variations. The weights are calculated using a tricubic or Gaussian weighting function (42,43), giving more importance to points closer to *x*_0_ in the fitting process.

#### Metrics

Quantifiable measurements are required to evaluate and contrast Ca^2+^ activity from control and perturbed or pathological experimental conditions (47,48). The established technique for detecting this activity involves using fluorescent Ca^2+^ indicators, which display changes in fluorescence intensity to indicate variations in Ca^2+^ levels. Parameters such as Ca^2+^ signal amplitude, area under the curve, and transient frequency can provide insights into concentration-response relationships for different stimuli (49). Analyzing these types of metrics helps to interpret Ca^2+^ responses across different experimental conditions and organisms, allowing to find the difference and similarities in cellular response. In the following section, we outline the metrics that can be extracted from multiple transients using CalciumInsights and how they could be interpreted in a biological context (Fig 1).

**Fig 1.**
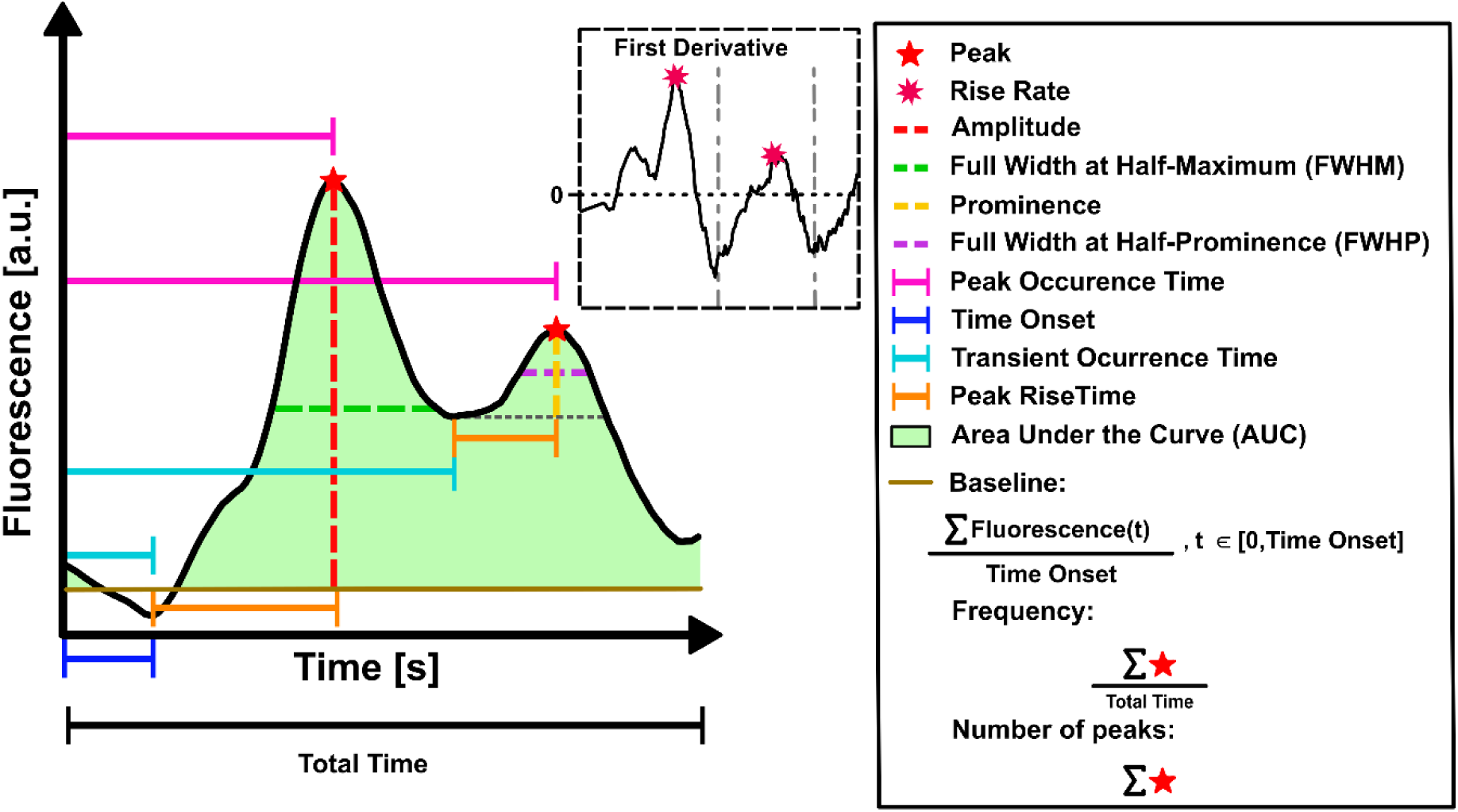
Ca^2+^ transients metrics: Representative metrics that can be extracted from the Ca^2+^ transients using CalciumInsights: Peak, Rise Rate, Amplitude, Full Width at Half-Maximum (FWHM), Prominence, Full Width at Half-Prominence (FWHP), Peak Occurrence Time, Time Onset, Transient Occurrence Time, Peak Rise Time, Baseline, Area Under the Curve (AUC), Number of Peaks, and Frequency.

##### Peak

The peak is the maximum fluorescence ratio for the duration of a transient (50).

##### Amplitude or Peak Height

The peak height is the difference between the baseline and peak, i.e., resting state and the amplitude of the transient respectively, (50,51) constructed using the following formula:

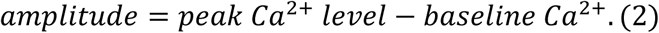

##### Full Width at Half-Maximum (FWHM)

Response duration is the time between half amplitude on the ascent and descent of the transient (48).

##### Prominence

Prominence refers to the distinctiveness of a peak in a Ca^2+^ signal. It measures how much a specific peak stands out from the surrounding fluctuations in Ca^2+^ concentration. Prominence considers both the height of the peak and its relative position compared to a neighboring transient event characterized by a temporary increase in Ca^2+^ concentration within a cell (47).

##### Full Width at Half-Prominence (FWHP)

The full width at half-prominence represents the width of a signal, typically a peak or an event, at the level where its prominence is equal to half of its maximum prominence. It measures the width of a feature in a signal at the point where its intensity has decreased by half from the peak value (47). This parameter is also referred to as transient duration 50 (TD50) (52).

##### Peak Occurrence Time

The peak occurrence time refers to the specific moment or point in time when a Ca^2+^ signal reaches its maximum amplitude or concentration during a transient event. This time provides information about when this increase reaches its zenith (53).

##### Time Onset

Time onset refers to the moment when the increase in Ca^2+^ concentration within a cell begins. It marks the initiation or starting point of the transient event (48,54).

##### Transient Occurrence Time

Transient occurrence time refers to the specific moment or time a transient event occurs. It signifies the initiation or onset of the transient event and is typically measured from the beginning of the recording or a specific reference point.

##### Peak Rise Time

Peak rise time is the time it takes a Ca^2+^ transient to peak from the baseline (55,56).

##### Baseline

The baseline is the fluorescence at the beginning of the transient (50); i.e., the initial level of Ca^2+^ before a signal occurs (56).

##### Area Under the Curve (AUC)

The area under the curve is a value obtained through numerical integration to quantify the amount of Ca^2+^ over a specific period (57).

##### Number of Peaks

The number of peaks denotes the number of Ca^2+^ transients or spikes in a signal. A peak is a pattern of abrupt change within the dataset, characterized by an ascent followed by a descent. This definition is implemented by analyzing the sign changes in the differences between adjacent data points in the dataset (58–60).

##### Frequency

Frequency represents how often an event repeats over time. It refers to the oscillatory feature of a Ca^2+^ signal, i.e., the number of peaks of Ca^2+^ divided by the total time of the signal (61).

##### Rise Rate

This metric is the maximum rate of change (first derivative) of the fluorescence during the transient (see inset in Fig 1). This expression can be used to quantify how quickly the average frequency is increasing in relation to time (62), and the maximal velocity is usually reported (63).

The specifications of how each parameter was calculated and implemented are provided in the user manual.

#### Filters

CalciumInsights provides filters that allow to fine-tune the peak detection algorithm. This helps the user to identify peaks that are relevant for biological system.

##### Minimum Peak Height

The minimum peak height is a predefined threshold that the transients must exceed to be considered significant. This threshold is established depending on the specific characteristics of the data and the analytical goals, and it helps distinguish relevant peaks from background noise or minor fluctuations.

##### Minimum Peak Distance

The minimum peak distance refers to the smallest allowable separation between consecutive transients. Specifying a minimum peak distance helps ensure that only distinct transients are considered. This parameter is essential for preventing the detection of closely spaced peaks as one single peak and avoiding false positives in peak identification.

##### Peak Ascent

The peak ascent refers to the minimum number of discrete time steps before reaching a peak (64).

##### Peak Descent

The peak descent refers to the minimum number of discrete time steps after the peak (64).

##### Minimum Width Threshold FWHP_min_

The minimum width threshold introduces a minimum width parameter that allows the user to set a threshold for the minimum allowable FWHP.

### Experimental Datasets

To assess the efficiency of CalciumInsights for analyzing Ca^2+^ traces collected from different organisms, we applied its workflow to extract all the metrics defined previously (Fig 1) from several sets of biological data. Data were derived from Ca^2+^ signal measurements in *Danio rerio, Arabidopsis thaliana,* and *Drosophila melanogaster,* and the experimental protocols can be found in the following section.

#### Danio rerio

The zebrafish experiment was conducted following internationally accepted standards. The Animal Care and Use Protocol was approved by The Purdue Animal Care and Use Committee (PACUC), adhering to the Guidelines for the Use of Zebrafish in the NIH Intramural Research Program (protocol number: 1401001018). Larvae were produced from *Tg(krt4:GAL4, UAS:GCaMP6f)* in a mixed AB/TL background that expresses the GCaMP6f (65) Ca^2+^ indicator in the outer epithelial layer (66). The larvae were raised in an E3 medium at 28°C until 3 days post fertilization (dpf) for experiments (67). Larvae were treated with 100 nM of MemGlow^TM^ 560 live membrane dye (Cytoskeleton, Cat #: MG02-10) for 20 min, anesthetized with 164 mg/L tricaine methanesulfonate (MedChemExpress, Cat. No.: HY-W011777), then immobilized in a custom-designed microfluidics device (68). The caudal tail fin fold was centered in the frame and imaged for 60 min at 1 frame per min on a Nikon A1Rmp with a Plan Fluor 20x 0.5 NA air objective on green (488-nm excitation, 510-nm emission) and red (561-nm excitation, 610-nm emission) channels, with a Z-stack spanning the full thickness of the fin. At the 60-min mark, imaging was paused, and latrunculin A (Cayman, Item No. 10010630) was added to achieve a final concentration of 200 µM. Imaging was then resumed for an additional 90 min. Fiji (19) was used to extract Ca^2+^ trace values from a subset of visible epidermal cells. Individual cells were segmented by hand utilizing the polyline ROI selection tool, followed by measurement of mean intensity within the ROI. Each ROI was manually shifted throughout the video to account for sample movement within the frame. Fluorescence changes were calculated as (F-F_0_)/F_0_, where F_0_ is the lowest mean fluorescence intensity demonstrated by that particular cell over the entire imaging period (Fig 5A, D).

#### Arabidopsis thaliana

*Arabidopsis thaliana* (Columbia-0 ecotype) expressing R-GECO1, and intensiometric Ca^2+^ reporter (69), was used in this work. Arabidopsis seeds were surface sterilized and stratified at 4°C for 3 days on half-strength Murashige and Skoog (MS) medium supplemented with 1% sucrose and 0.8% agar. Plants were grown under long-day lighting conditions (16 h light/8 h dark) at 21°C for 7 days. Before imagining, cotyledons were excised from seedlings and soaked in water overnight. The cotyledons were then mounted in custom-made chambers for bottom imaging as described previously with modifications (69). Cotyledons were placed in water at the bottom of the chamber with a cotton pad on top as a spacer, and then thin agar blocks were placed over the cotton to stabilize it. Samples were allowed to rest in the chamber for at least 40 min before imaging.

Epidermal cells from the apical region of the abaxial side of cotyledons were imaged for R-GECO1 fluorescence. Spinning-disk confocal microscopy (SDCM) was performed using a Yokogawa scanner unit CSU-X1-A1 mounted on an Olympus IX-83 microscope, equipped with a 20X 0.4–numerical aperture (NA) Plan C Achromat Dry objective (Olympus) and an Andor iXon Ultra 897BV EMCCD camera (Andor Technology). R-GECO1 fluorescence was excited with the 561-nm laser line, and emission was collected through a 610/37-nm filter. Time-lapse image series were collected at a single focal plane with 5-s intervals for 481 frames (40 min). Image acquisition was paused after the 120th frame. A two-fold concentration of the immunogenic peptide flg22 (GenScript) was added to the imaging chamber to achieve a final concentration of 1 µM (Fig 5B, E). Fiji (19) was used to extract Ca^2+^ fluorescence intensity values from individual epidermal cells. The puzzle piece-shaped epidermal cells were manually traced using the freehand line tool, and the line selections with a pixel width of 2 were used as ROIs to extract the raw intensity values. The fluorescence changes were calculated as (F-F_0_)/F_0_ or ΔF/F_0_, where F_0_ is the average intensity of a stable period in the first 120 frames after subtracting the background.

#### Drosophila melanogaster

Fly stocks and culture were grown and maintained at 25 °C with a twelve-hour light cycle. The recombined Ca^2+^ sensor line developed at the Zartman Lab (using the Bloomington Drosophila Stock Center fly lines #2575 and #42747) has the genotype nubbin-GAL4, UAS-GCaMP6f/CyO. Flies were grown in a 15:5 ratio of female to male and staged for 6 hours to ensure the collection of wandering third instar larvae. Dissected wing imaginal discs were cultured with organ culture media with 1 mM Yoda1 or insulin to stimulate Ca^2+^ dynamics. Dissected discs were mounted in polyethylene terephthalate laminate (PETL) microfluidic devices where cultured media was introduced for long-term imaging (70). The devices were then imaged on a confocal microscope, and culture media was introduced using syringe pumps (Harvard Apparatus). The organ culture media consists of Grace’s insect medium supplemented with Bis-Tris, fetal bovine serum, penicillin and streptomycin, and 20-Hydroxyecdysone (71). Initial time points were taken as a control in Grace’s media without any drug perturbation.

Imaging was acquired using a Nikon Eclipse Ti confocal microscope with a Yokogawa spinning disc. Image data were collected on an iXonEM+cooled CCD camera (Andor Technology, South Windsor, CT) using MetaMorph v7.7.9 software (Molecular Devices, Sunnyvale, CA). Discs were imaged throughout the entire depth with a step size of 1 µm, depending on sample thickness, with a 40x or 60x oil objective with 200-ms exposure time, 50 nW, and 488-nm laser exposure at 44% laser power. The imaging was performed from the apical to the basal surface so that periodical cells were imaged first, followed by the columnar cells of the wing disc. Post-processing with Fiji (19) was used to adjust the brightness and contrast of the imaging data and create video files of raw data (Fig 5C, F). Fiji was used to extract Ca^2+^ fluorescence intensity values from individual epithelial cells of the wing disc. Each spiking cell was manually traced using the rounded square tool as ROIs to extract the raw intensity values. The fluorescence changes were calculated for all 360 frames using the Plot Z-axis Profile tool in Fiji.

#### Designing CalciumInsights

R is an open-source software and a widely used programming language for statistical computation and data analysis (72). Shiny is a framework for creating dynamic and interactive web applications that fully harnesses all features of R (40). Compared to other programming languages, it stands out for its significant contribution to developing numerous web applications (72). We used RStudio (1.3.1093) and R (v4.0.3) to develop a web application. This application has been modularized using the Golem package (39), enabling the bundling of two separate modules. Furthermore, it leverages the Shiny framework to provide a GUI. The first module aims to enhance Ca^2+^ signals by removing noise using the Loess algorithm. The second module focuses on data analysis, providing a comprehensive characterization of Ca^2+^ signals. We utilized several R functions including findPeaks (64), integrate (73), ggplot (74), and Loess (75). Additionally, we incorporated custom functions, such as Rise Rate, Full Width at Half-Maximum (FWHM), Prominence, FWHP, Time Onset, Transient Occurrence Time, Peak Rise Time, Baseline, and Frequency, which were designed to calculate specific metrics from Ca^2+^ signals.

## Results

### General Overview of CalciumInsights

The workflow of CalciumInsights focuses on simplicity without sacrificing analytical power (Fig 2). Input files accepted by CalciumInsights, include .csv or. tsv file format and must contain Ca^2+^ fluorescence intensity values (in raw or normalized form) for various annotated ROIs and timestamps for each frame. These datasets can be extracted from open-source image processing packages like Fiji. The analysis appears in the graphical interface as soon as the document is input into the workflow (Fig 2). As shown in Fig 3, CalciumInsights provides a wide range of metrics extracted from the transients, facilitating real-time visualization of the study. Please refer to the user manual in the supplementary material for detailed instructions on effectively utilizing the graphical interface.

**Fig 2.**
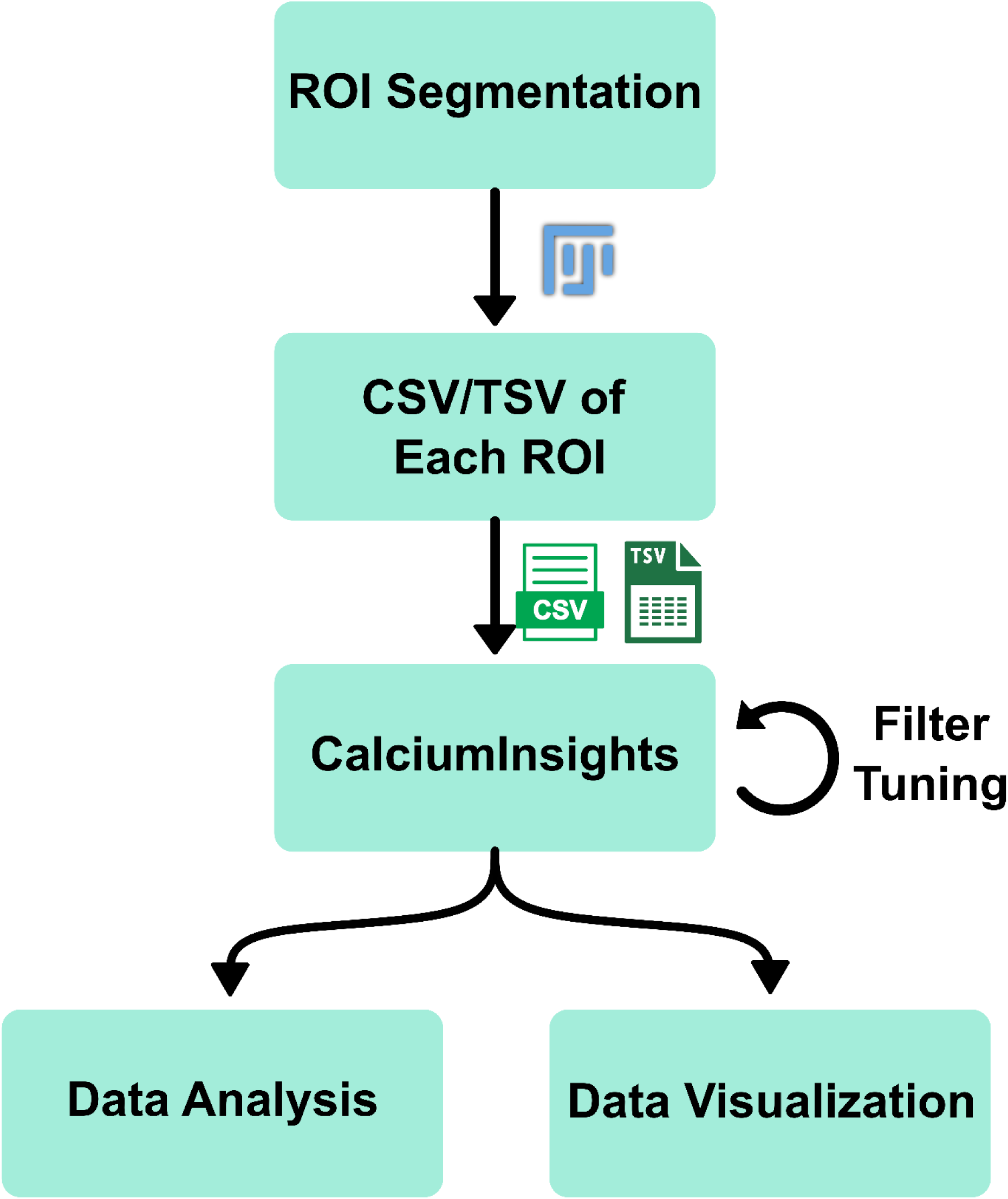
CalciumInsights workflow. The input requires a data frame encoded in a .csv or .tsv file. Then, the user enters the iterative process of fine-tuning filters. After fine-tuning, metrics and visualizations can be downloaded for further analysis. See also Fig. S1 for a more in-depth representation.

**Fig 3.**
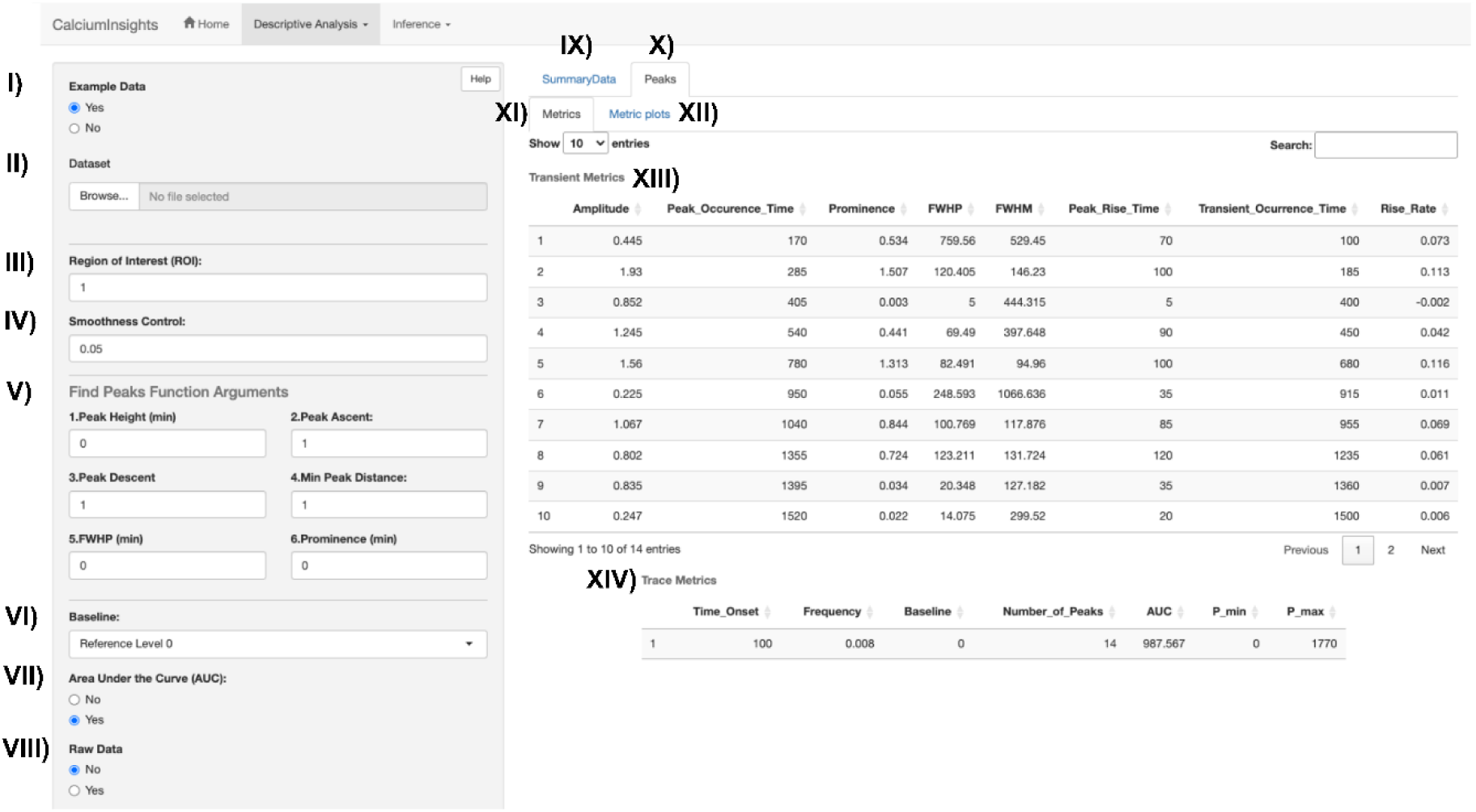
Menu option to fine-tune parameters and results of estimated metrics. The panel on the left provides options to import example data (I), import user’s data (II), select the ROI to analyze (III), define Smoothness Control (IV), and adjust the filters for the Find Peaks function (V), define the baseline that represents the signal (VI), plot the area under the curve with the smoothed signal (VII) and plot the raw signal together with the smoothed signal (VIII). The controls define the page on the right. A summary of the ROI’s metrics can be selected (IX) or the peaks of a specific ROI (X). In the latter, the metrics can be seen as the numerical values (XI) or in a plot (XII). A table is provided with the numerical value of the metrics of each transient (XIII) and the complete trace (XIV).

Users can adjust the “span” parameter (also known as the “Smoothing Control” parameter) in CalciumInsights between values of 0 and 1 (Fig 3). For further information, please refer to the Methods section. CalciumInsights enables users to customize the analysis for any biological system they investigate by providing different filters.

Users can define and construct additional modules to add their metrics and visualizations to CalciumInsights using Golem. This enables the expansion of the tool’s current features and increases the options for customized analysis. Please refer to the user manual in the Supplementary Material for a more detailed description of defining various metrics and visualizations and how the tool integrates them.

Linear regression is conducted to fully exploit the Loess smoothing, with the smoothed fluorescence data after passing through the Loess filter serving as the dependent variable and the raw fluorescence data as the independent variable (Fig 4). This example illustrates how well the original raw data computed using the least square approach fits into the smoothed data model.

**Fig 4.**
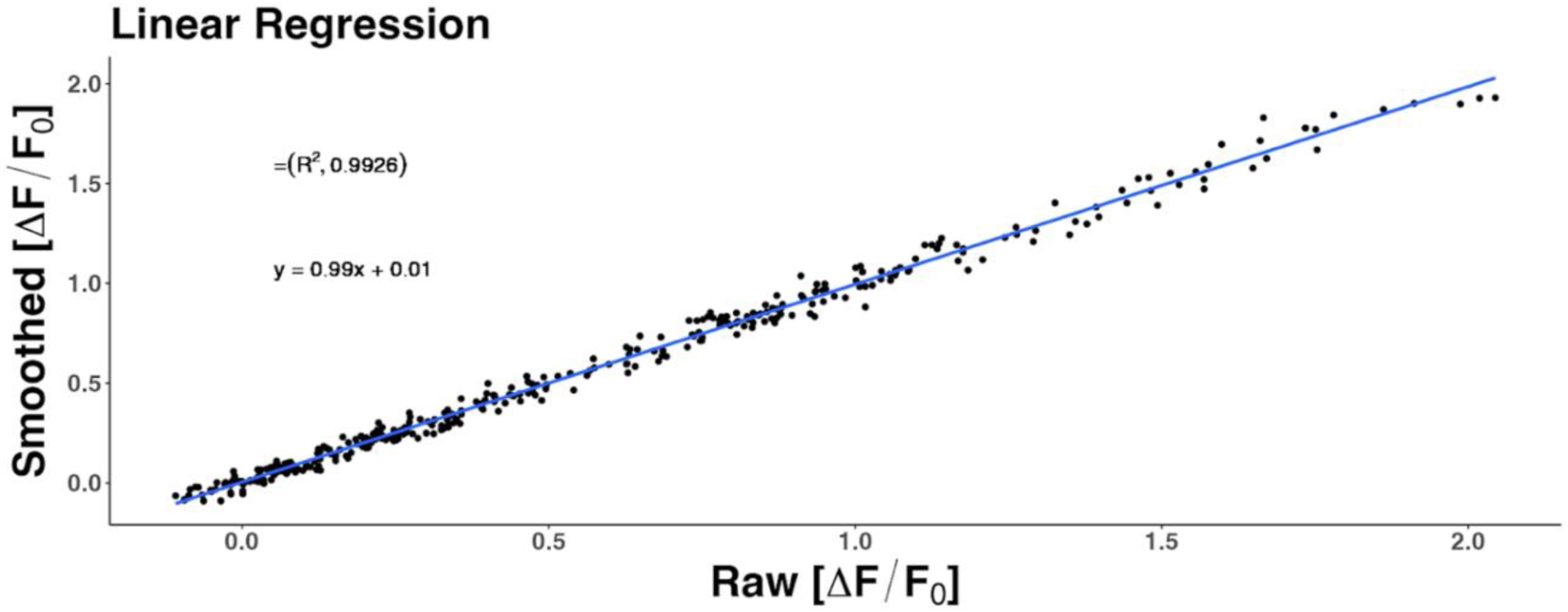
Linear regression between raw data and smoothed data. A linear regression between raw and smoothed datasets is generated to assess the smoothing via Loess. A better fit is indicated when *R*^2^ is closer to 1.

#### Testing

We assessed CalciumInsights using Ca^2+^ recordings obtained from the tissues and organisms depicted in Fig 5, which include the outer epithelial layer of the resting embryonic tail fin of *Danio rerio*, three days post-fertilization (Fig 5A), the epidermal cells from the cotyledon of *Arabidopsis thaliana* (Fig 5B), and an ex vivo wing imaginal disc from a third instar larva of *Drosophila melanogaster* (Fig 5C). Individual cells or regions exhibiting significant Ca^2+^ dynamics were identified as ROIs and manually annotated using Fiji (Fig 5).

**Fig 5.**
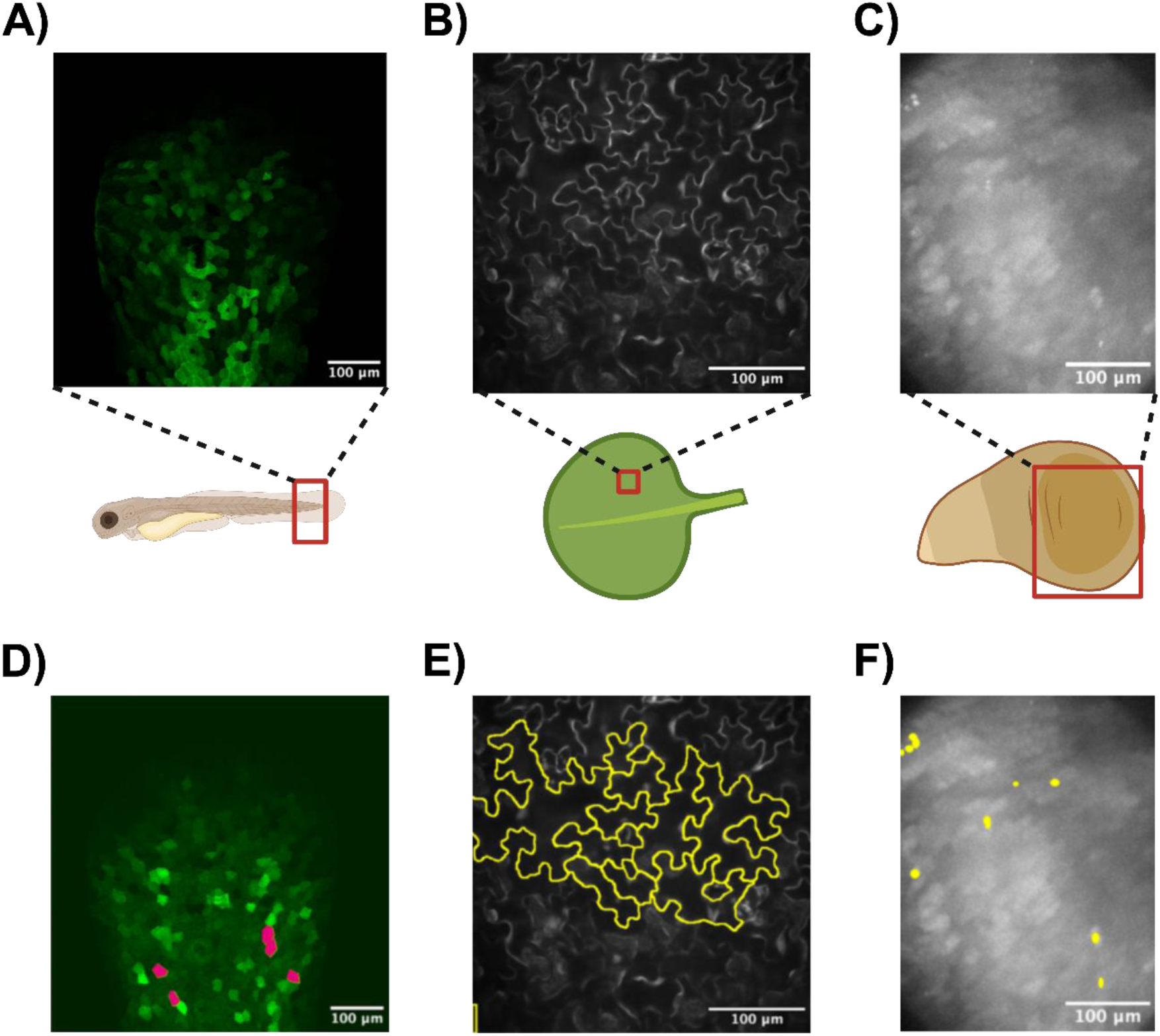
Calcium imaging of different biological systems and manual segmentation. A) Ca^2+^ signal in the outer epithelial layer of the resting *Danio rerio* tail fin at 3 days post fertilization (dpf) after the application of 200 µM latrunculin A. B) Ca^2+^ imaging of the cotyledon epidermal cells of *Arabidopsis thaliana* expressing R-GECO1 after application of 1 µM of the immunogenic peptide, flg22. C) Ca^2+^ signal in an *ex vivo* wing imaginal disc from a third instar larva of *Drosophila melanogaster* after applying 1 mM Yoda1 or insulin. D) the outer epithelial layer of the resting *Danio rerio* tail fin at 3 dpf, E) the cotyledon epidermal cells of *Arabidopsis thaliana*, and F) an *ex vivo* wing imaginal disc from a third instar larva of *Drosophila melanogaster*. Image created using BioRender.com.

Representative Ca^2+^ traces from the dataset are visualized using datasets available in Supplementary File S1, corresponding to the embryonic tail fin of the zebrafish, the epidermal cells from the cotyledon of *Arabidopsis thaliana*, and the ex vivo wing imaginal discs from a third instar larva of *Drosophila melanogaster*, respectively (Fig 6A, 7A, 8A). Box plots represent the distribution of the datasets, including the median, quartiles, and potential outliers. For each case, the application filters were carefully adjusted (Fig 6B, 7B, 8B).

**Fig 6.**
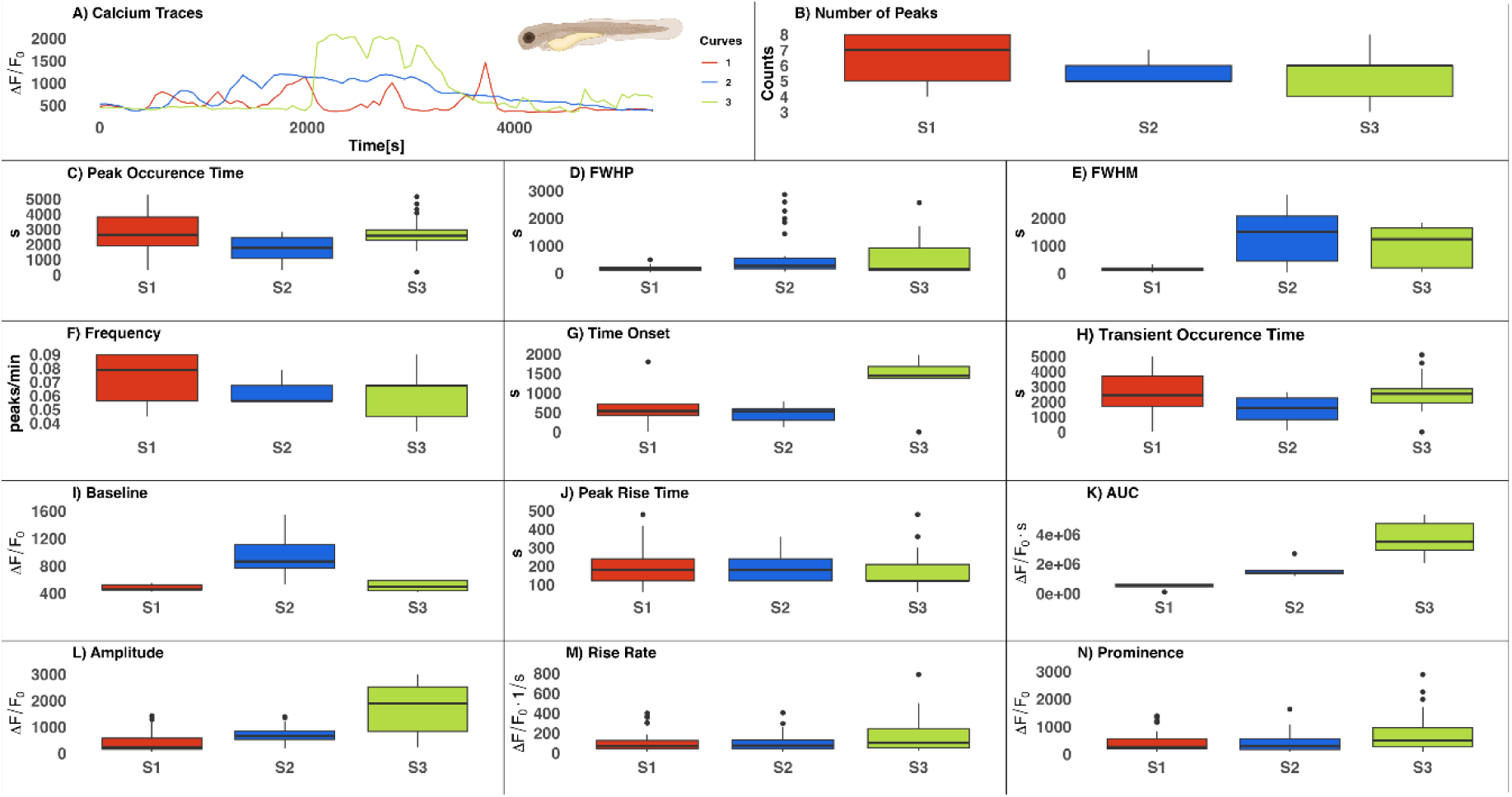
Calcium traces and metrics of the epithelial layer of the resting *Danio rerio* embryonic tail fin. A) Representative Ca^2+^ Traces (one per trial to show the shape), B) Number of Peaks, C) Peak Occurrence Time, D) FWHP, E) FWHM, F) Frequency, G) Time Onset, H) Transient Occurrence Time, I) Baseline, J) Peak Rise Time, K) AUC, L) Amplitude, M) Rise Rate, and N) Prominence. Smoothness Control: 0.05, Peak Height (min): 500, FWHP (min): 40, Prominence (min): 70, and Baseline: Min. The number of ROIs per biological replicate is 5, and data from 3 biological replicates (S1, S2, S3) are shown.

**Fig 7.**
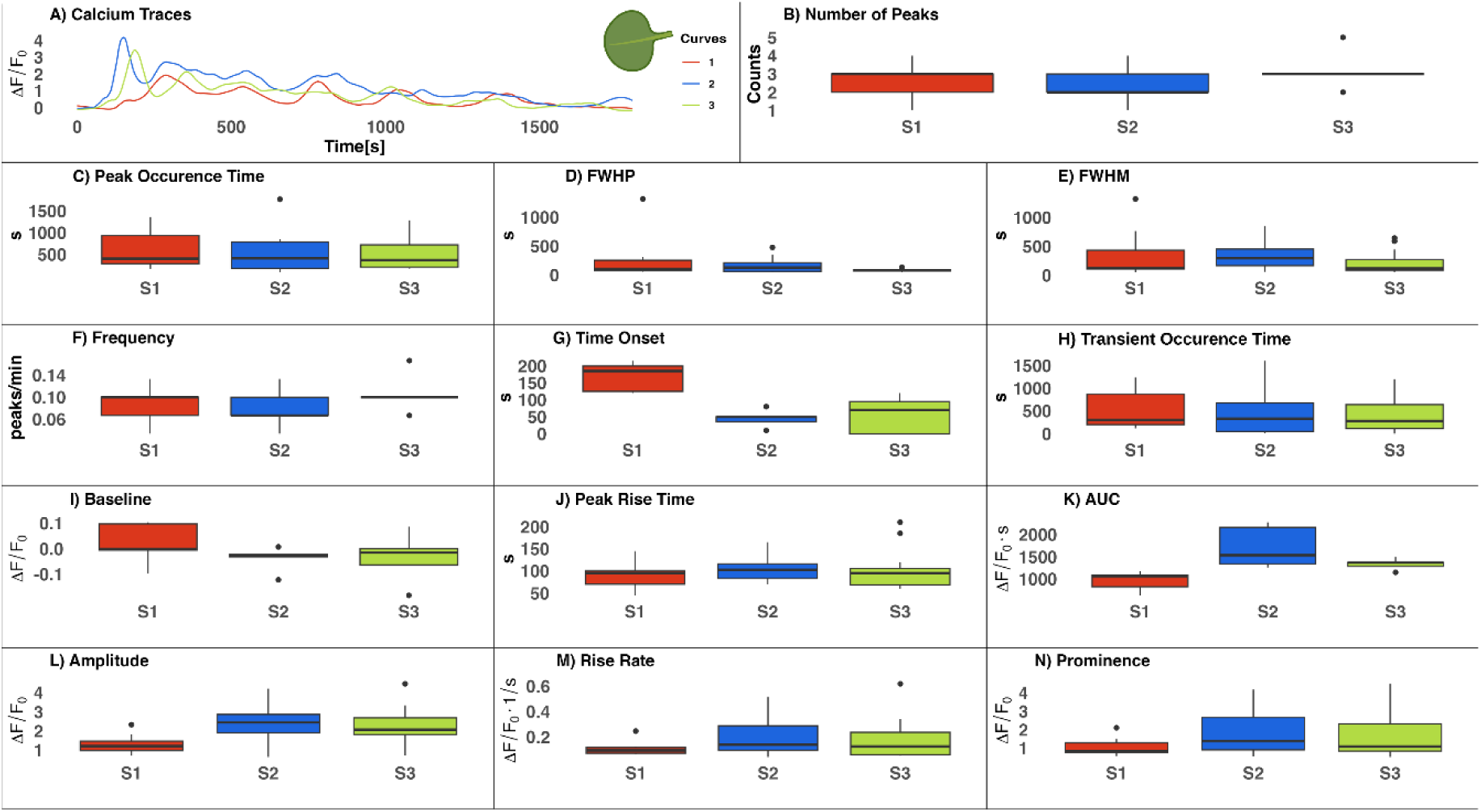
Calcium traces and metrics for the cotyledon epidermal cells from *Arabidopsis thaliana*. A) Representative Ca^2+^ Traces (one per biological replicate to show the shape), B) Number of Peaks, C) Peak Occurrence Time, D) FWHP, E) FWHM, F) Frequency, G) Time Onset, H) Transient Occurrence Time, I) Baseline, J) Peak Rise Time, K) AUC, L) Amplitude, M) Rise Rate, and N) Prominence. Smoothness Control: 0.05, Peak Height (min): 0.5, FWHP (min): 40, Prominence (min): 0.5, and baseline: standard definition. The number of ROIs per biological replicate (cotyledon) is equal to 5, and there are 3 biological replicates (S1, S2, S3).

**Fig 8.**
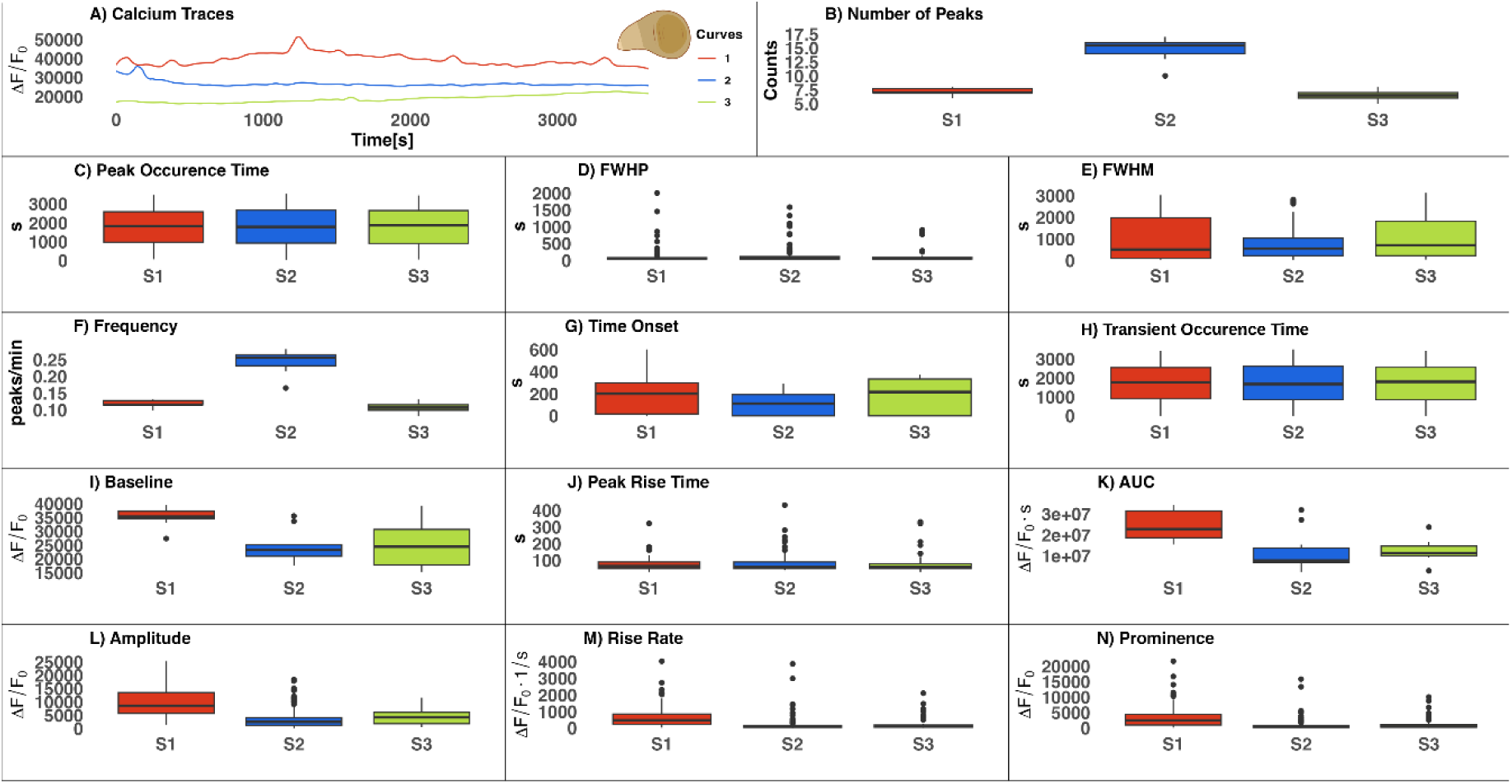
Calcium transients of *ex vivo* wing imaginal discs from a third instar larva of *Drosophila melanogaster.* A) Representative Ca^2+^ Traces (one per trial to show the shape), B) Number of Peaks, C) Peak Occurrence Time, D) FWHP, E) FWHM, F) Frequency, G) Time Onset, H) Transient Occurrence Time, I) Baseline, J) Peak Rise Time, K) AUC, L) Amplitude, M) Rise Rate, and N) Prominence. Smoothness Control: 0.04, Peak Height (min): 17000, Peak Ascend: 3, Peak Descend: 3, Min Peak Distance: 36, and baseline: standard definition. The number of ROIs per biological replicate is equal to 10, and 3 biological replicates (S1, S2, S3).

## Discussion

Given that Ca^2+^ dynamics influence cell fate, organ development, and response to the environment, its intricate role in cellular and tissue processes has long been appreciated (76,77). This study addresses the critical need for a comprehensive post-processing tool to analyze Ca^2+^ traces in cells, tissues, or organs and quantifiably characterize Ca^2+^ dynamics under normal, pathological, or inducible conditions. There is a lack of available post-processing tools for this purpose, which prompted the creation of new solutions (22–26). In this work, we present a new tool for post-processing analysis of Ca^2+^ dynamics in tissues and individual cells (Fig 1, 2). Among the many features of this program is a set of crucial metrics specifically tailored for Ca^2+^ imaging data analysis. Due to its flexibility and adaptability, it can be utilized across various biological systems and conditions. Additionally, its open-source architecture enables users to incorporate additional metrics and analyses to enhance its capabilities (30–33,35,36). The flexibility provided by this tool ensures its ability to adapt to changing research needs. By emphasizing user-friendliness, expandability, and ease of use for researchers without programming experience and fostering a collaborative environment within the scientific community, CalciumInsights aims to empower researchers from diverse backgrounds by expanding access to Ca^2+^ analysis tools.

The way researchers present Ca^2+^ signaling data can vary according to their objectives and audience. For example, if the primary aim is to illustrate patterns or changes over time, researchers may choose to display Ca^2+^ traces alongside corresponding statistical analyses. Alternatively, parameters such as the proportion of cells exhibiting detectable signals, signal amplitude, and other relevant metrics can be emphasized for a more thorough characterization of cellular or tissue responses to stimulation. This study presents changes in Ca^2+^ activity over time, along with multiple parameters from three different biological systems analyzed using CalciumInsights, offering a comprehensive characterization of cellular and tissue responses to stimulation (Fig 6-8). Subsequently, these metrics can facilitate establishing dose-response relationships for various stimuli.

In this study, Loess and Savitsky-Golay (Defined in the Supporting Information) smoothing were the two main techniques examined. Loess employs a non-parametric method to fit distinct polynomials to various data segments to capture intricate, nonlinear relationships. Unlike Savitzky-Golay, Loess does not assume that there is an underlying structure. Due to its weighted scheme, deliberately reducing the impact of isolated data points during the smoothing phase, Loess is also less susceptible to outliers. The ease of implementation also contributes to the popularity of Loess, with several software packages offering user-friendly tools for applying this method to different datasets (78). In this regard, Loess had a specific advantage over Savitzky-Golay, as the latter has the limitation of only smoothing or computing a derivative for a subset of the analyzed data; specifically, the data points at the beginning and end of a trace are not smoothed but instead removed. The explanation lies in the window’s symmetrical requirement around the smoothing points. The polynomial used for regression around each point depends on a minimum window size and consumes too many points, rendering it impractical for most evaluated metrics (79). The only exception is the Rise Rate, which refers to the maximum derivative within the interval when a transient begins to rise until it reaches its corresponding peak. Savitzky-Golay was used to compute this specific metric after the original smoothing.

We evaluated whether the Loess smoothing technique impacts the calculation of the metrics by comparing the values extracted from the same trace with and without smoothing (Fig S5). The results showed no statistically significant differences (p-value > 0.05). In cases where the p-value was closer to 0.05, specifically for Peak Rise Time and Full Width at Half Peak (FWHP), these differences can be attributed to the metrics’ dependence on the derivative of the trace (Table S1). Calculating this derivative using numerical methods is highly sensitive to noise, making these metrics more susceptible to noise-related variability than others.

The versatility and adaptability of CalciumInsights were demonstrated when applied to three different model organisms exhibiting Ca^2+^ dynamics (Fig 5). Our tool effectively deciphered the parameters of Ca^2+^ signals in various cellular structures, including the multi-edged polygonal cells of the zebrafish caudal fin, the puzzle-piece-shaped epidermal cells from a plant cotyledon, and the epithelial cells of the fruit fly (Fig 5). Various images taken at different time intervals for these biological samples were successfully analyzed, ranging from 54 to 361 frames with intervals of 5 to 60 seconds (SI Metadata). This highlights CalciumInsights’ ability to measure and calculate a wide range of parameter values accurately. For instance, these three specimens’ AUC values varied from 2000 to 4×10^6^ (Fig 6–8). Overall, our tool demonstrates the ability to analyze cellular Ca^2+^ signaling parameters regardless of the biological model of origin, even across different kingdoms.

The current version of CalciumInsights requires the user to generate time trace data from ROIs using a separate tool. This capability allows the analysis of various cell and tissue types, regardless of origin. The flexibility and extensibility of CalciumInsights also support the implementation of additional modules or metrics, enabling users to incorporate methods specific to their system.

Further, programming expertise is required to integrate new metrics into the platform, which may limit accessibility for non-computational bench researchers. Striking a balance between advanced functionality and accessibility is a challenge in the continuous development of these technologies. To this end, we provide an instruction manual for utilizing our tool and an application integration template for adding new features and modules (see Supporting Information).

Optimization methods can be integrated to adjust the range of the Loess smoothing technique in future updates of CalciumInsights, functional decomposition, Fourier Transform analysis, and analysis of individual transients. As an enhancement, statistical tests such as functional ANOVA can also be added to CalciumInsight’s workflow to enable statistical comparisons among diverse samples subjected to different stimuli. Thanks to the software’s flexibility, all these improvements are feasible, with the limitation that the enhancements must be made exclusively in the R language (80).

In conclusion, this study introduces a tool for Ca^2+^ signature analysis in tissues and cells, facilitating comparisons among different biological systems through a unique integrated analysis platform. By developing an accessible and adaptable tool capable of evolving with changing perspectives in Ca^2+^ signal analysis, CalciumInsights will continue shedding light on the role of Ca^2+^ in vital processes and enable biological research breakthroughs.

## Author Contributions

Conceptualization: ABT, CJS, CEIB, DG, DU, DSG, GIMC, GR, JZ, MCR, MSM, NCPR, QD, SRCB, SGT, WZ. Data curation: DSG, GIMC, MSM, NCPR, SRCB, ST, WZ. Formal analysis: CJS, CEIB, DG, DU, DSG, GIMC, GR, JZ, MCR, MSM, NCPR, QD, SRCB, SGT, WZ. Funding acquisition: ABT, CJS, CEIB, DU, JZ, MCR, QD. Investigation: CJS, CEIB, DG, DU, DSG, GIMC, GR, JZ, MCR, MSM, NCPR, QD, SRCB, SGT, WZ. Methodology: CJS, CEIB, GZ, DSG, GIMC, GR, JZ, MCR, MSM, NCPR, QD, SRCB, SGT, WZ. Project administration: DSG, GIMC, NCPR, SRCB. Resources: ABT, CJS, CEIB, DU, GR, JZ, MCR, QD. Software: DSG, GIMC, NCPR, SRCB, WZ. Supervision: ABT, CJS, CEIB, DU, GR, JZ, MCR, QD. Validation: CJS, CEIB, DG, DU, DSG, GIMC, GR, JZ, MCR, MSM, NCPR, QD, SRCB, SGT, WZ. Visualization: DG, DSG, GIMC, GR, JZ, MCR, MSM, NCPR, QD, SRCB, SGT, WZ. Writing Original Draft preparation: DG, DSG, GIMC, MSM, NCPR, SRCB, ST, WZ. Writing review-editing: CJS, CEIB, DG, DU, DSG, GIMC, GR, JZ, MCR, MSM, NCPR, QD, SRCB, SGT, WZ.

## Acknowledgments

We want to acknowledge Blanca Maria Romero Dussaillant and Augustin Luna for their computational insights and Agustín Guerrero Hernández for his advice about the analysis of Ca^2+^ traces. This work is based upon efforts supported by the EMBRIO Institute, NSF contract #2120200, a National Science Foundation (NSF) Biology Integration Institute. GIMC would like to acknowledge the Neale Silva Scholarship of the University of Wisconsin-Madison and Dr. Ophelia Venturelli, who helped with the financial support during the development of this project. SRCB would like to thank the Department of Mathematics at the University of Puerto Rico at Mayagüez for their financial support.

## Consent for publication

All authors have reviewed the manuscript and approved the final draft for publication.

## Resource availability

Lead contact: Further information and requests for data may be directed to and will be fulfilled by Mauricio Cabrera

Code: All codes used are publicly available in GitHub at https://github.com/AOG-Lab/CalciumInsights

## References

1. Berridge MJ, Lipp P, Bootman MD. The versatility and universality of calcium signalling. Nat Rev Mol Cell Biol [Internet]. 2000 Oct [cited 2024 Oct 21];1(1):11–21. Available from: https://www.nature.com/articles/35036035

2. Sanderson MJ, Charles AC, Boitano S, Dirksen ER. Mechanisms and function of intercellular calcium signaling. Mol Cell Endocrinol. 1994 Jan;98(2):173–87.

3. Webb SE, Miller AL. Calcium signalling during embryonic development. Nat Rev Mol Cell Biol. 2003 Jul;4(7):539–51.

4. Luan S, Wang C. Calcium Signaling Mechanisms Across Kingdoms. Annu Rev Cell Dev Biol. 2021 Oct 6;37:311–40.

5. Dodd AN, Kudla J, Sanders D. The language of calcium signaling. Annu Rev Plant Biol. 2010;61:593–620.

6. Berridge MJ, Bootman MD, Roderick HL. Calcium signalling: dynamics, homeostasis and remodelling. Nat Rev Mol Cell Biol [Internet]. 2003 Jul [cited 2024 Oct 21];4(7):517–29. Available from: https://www.nature.com/articles/nrm1155

7. Stewart TA, Davis FM. An element for development: Calcium signaling in mammalian reproduction and development. Biochim Biophys Acta BBA - Mol Cell Res [Internet]. 2019 Jul 1 [cited 2024 Oct 21];1866(7):1230–8. Available from: https://www.sciencedirect.com/science/article/pii/S0167488919300230

8. Islam MdS, editor. Calcium Signaling [Internet]. Cham: Springer International Publishing; 2020 [cited 2024 Oct 21]. (Advances in Experimental Medicine and Biology; vol. 1131). Available from: http://link.springer.com/10.1007/978-3-030-12457-1

9. Brodskiy PA, Wu Q, Soundarrajan DK, Huizar FJ, Chen J, Liang P, et al. Decoding Calcium Signaling Dynamics during *Drosophila* Wing Disc Development. Biophys J [Internet]. 2019 Feb 19 [cited 2024 Oct 21];116(4):725–40. Available from: https://www.sciencedirect.com/science/article/pii/S0006349519300232

10. Sipka T, Peroceschi R, Hassan-Abdi R, Groß M, Ellett F, Begon-Pescia C, et al. Damage-Induced Calcium Signaling and Reactive Oxygen Species Mediate Macrophage Activation in Zebrafish. Front Immunol. 2021;12:636585.

11. Soundarrajan DK, Huizar FJ, Paravitorghabeh R, Robinett T, Zartman JJ. From spikes to intercellular waves: Tuning intercellular calcium signaling dynamics modulates organ size control. PLoS Comput Biol [Internet]. 2021 Nov 1 [cited 2024 Oct 21];17(11):e1009543. Available from: https://pmc.ncbi.nlm.nih.gov/articles/PMC8601605/

12. Foskett JK, White C, Cheung KH, Mak D on D. Inositol Trisphosphate Receptor Ca2+ Release Channels. Physiol Rev [Internet]. 2007 Apr [cited 2024 Oct 21];87(2):593. Available from: https://pmc.ncbi.nlm.nih.gov/articles/PMC2901638/

13. Webb SE, Miller AL. Aequorin-based genetic approaches to visualize Ca2 + signaling in developing animal systems. Biochim Biophys Acta BBA - Gen Subj [Internet]. 2012 Aug 1 [cited 2024 Oct 21];1820(8):1160–8. Available from: https://www.sciencedirect.com/science/article/pii/S030441651100300X

14. Paredes RM, Etzler JC, Watts LT, Zheng W, Lechleiter JD. Chemical calcium indicators. Methods San Diego Calif. 2008 Nov;46(3):143–51.

15. Mollinedo-Gajate I, Song C, Knöpfel T. Genetically Encoded Fluorescent Calcium and Voltage Indicators. In: Barrett JE, Page CP, Michel MC, editors. Concepts and Principles of Pharmacology: 100 Years of the Handbook of Experimental Pharmacology [Internet]. Cham: Springer International Publishing; 2019 [cited 2024 Oct 21]. p. 209–29. Available from: 10.1007/164_2019_299

16. Li ES, Saha MS. Optimizing Calcium Detection Methods in Animal Systems: A Sandbox for Synthetic Biology. Biomolecules [Internet]. 2021 Feb 24 [cited 2024 Oct 21];11(3):343. Available from: https://pmc.ncbi.nlm.nih.gov/articles/PMC7996158/

17. Redolfi N, García-Casas P, Fornetto C, Sonda S, Pizzo P, Pendin D. Lighting Up Ca2+ Dynamics in Animal Models. Cells [Internet]. 2021 Aug [cited 2024 Oct 21];10(8):2133. Available from: https://www.mdpi.com/2073-4409/10/8/2133

18. Shipley FB, Clark CM, Alkema MJ, Leifer AM. Simultaneous optogenetic manipulation and calcium imaging in freely moving C. elegans. Front Neural Circuits. 2014;8:28.

19. Schindelin J, Arganda-Carreras I, Frise E, Kaynig V, Longair M, Pietzsch T, et al. Fiji: an open-source platform for biological-image analysis. Nat Methods [Internet]. 2012 Jul [cited 2024 Oct 21];9(7):676–82. Available from: https://www.nature.com/articles/nmeth.2019

20. Stringer C, Wang T, Michaelos M, Pachitariu M. Cellpose: a generalist algorithm for cellular segmentation. Nat Methods. 2021 Jan;18(1):100–6.

21. Giovannucci A, Friedrich J, Gunn P, Kalfon J, Brown BL, Koay SA, et al. CaImAn an open source tool for scalable calcium imaging data analysis. eLife. 2019 Jan 17;8:e38173.

22. Pasqualin C, Gannier F, Yu A, Benoist D, Findlay I, Bordy R, et al. Spiky: An ImageJ Plugin for Data Analysis of Functional Cardiac and Cardiomyocyte Studies. J Imaging. 2022 Apr 1;8(4):95.

23. Psaras Y, Margara F, Cicconet M, Sparrow AJ, Repetti GG, Schmid M, et al. CalTrack: High-Throughput Automated Calcium Transient Analysis in Cardiomyocytes. Circ Res. 2021 Jul 9;129(2):326–41.

24. Mukamel EA, Nimmerjahn A, Schnitzer MJ. Automated analysis of cellular signals from large-scale calcium imaging data. Neuron. 2009 Sep 24;63(6):747–60.

25. Velez Rueda AJ, Gonano LA, Smith AG, Parisi G, Fornasari MS, Sommese LM. CardIAP: calcium transients confocal image analysis tool. Front Bioinforma [Internet]. 2023 Jul 14 [cited 2024 Oct 21];3. Available from: https://www.frontiersin.org/journals/bioinformatics/articles/10.3389/fbinf.2023.1137815/full

26. Francis M, Qian X, Charbel C, Ledoux J, Parker JC, Taylor MS. Automated region of interest analysis of dynamic Ca^2^+ signals in image sequences. Am J Physiol Cell Physiol. 2012 Aug 1;303(3):C236–243.

27. Kumaraswamy A, Raiser G, Galizia CG. PyView: A general purpose tool for analyzing calcium imaging data. J Open Source Softw [Internet]. 2023 Feb 21 [cited 2025 Mar 20];8(82):4936. Available from: https://joss.theoj.org/papers/10.21105/joss.04936

28. Coleman P, Hogg PW, Toth TD, Haas K. PyNeuroTrace - Python code for neural activity time series. J Open Source Softw [Internet]. 2024 Aug 15 [cited 2025 Mar 20];9(100):6877. Available from: https://joss.theoj.org/papers/10.21105/joss.06877

29. Radstake FDW, Raaijmakers EAL, Luttge R, Zinger S, Frimat JP. CALIMA: The semi-automated open-source calcium imaging analyzer. Comput Methods Programs Biomed [Internet]. 2019 Oct [cited 2025 Mar 20];179:104991. Available from: https://www.ncbi.nlm.nih.gov/pmc/articles/PMC6718774/

30. Webmaster ISC. CytoSolver Transient Analysis Tool [Internet]. IonOptix. [cited 2024 Oct 21]. Available from: https://www.ionoptix.com/products/software/cytosolver-transient-analysis-tool/

31. pCLAMP, Clampfit: Perform a typical single-channel analysis usingthe Clampfit Software [Internet]. [cited 2024 Oct 21]. Available from: https://support.moleculardevices.com/s/article/pCLAMP-Clampfit-Perform-a-typical-single-channel-analysis-usingthe-Clampfit-Software

32. Igor Pro® | Igor Pro by WaveMetrics [Internet]. [cited 2024 Oct 21]. Available from: https://www.wavemetrics.com/products/igorpro

33. Gregory R. Vala Sciences. [cited 2024 Oct 21]. CyteSeer. Available from: https://valasciences.com/cyteseer/

34. Cheng H, Song LS, Shirokova N, González A, Lakatta EG, Ríos E, et al. Amplitude distribution of calcium sparks in confocal images: theory and studies with an automatic detection method. Biophys J [Internet]. 1999 Feb [cited 2024 Oct 21];76(2):606. Available from: https://pmc.ncbi.nlm.nih.gov/articles/PMC1300067/

35. Krzesiak A, Cognard C, Sebille S, Carré G, Bosquet L, Delpech N. High-intensity intermittent training is as effective as moderate continuous training, and not deleterious, in cardiomyocyte remodeling of hypertensive rats. J Appl Physiol Bethesda Md 1985. 2019 Apr 1;126(4):903–15.

36. Sebille S, Cantereau A, Vandebrouck C, Balghi H, Constantin B, Raymond G, et al. Calcium sparks in muscle cells: interactive procedures for automatic detection and measurements on line-scan confocal images series. Comput Methods Programs Biomed [Internet]. 2005 Jan 1 [cited 2024 Oct 21];77(1):57–70. Available from: https://www.sciencedirect.com/science/article/pii/S0169260704001646

37. Pearce JM. Sponsored Libre Research Agreements to Create Free and Open Source Software and Hardware. Inventions [Internet]. 2018 Sep [cited 2024 Oct 21];3(3):44. Available from: https://www.mdpi.com/2411-5134/3/3/44

38. Wickham H. Welcome | Mastering Shiny [Internet]. [cited 2024 Oct 21]. Available from: https://mastering-shiny.org/

39. ThinkR-open/golem [Internet]. ThinkR; 2024 [cited 2024 Oct 21]. Available from: https://github.com/ThinkR-open/golem

40. Jia L, Yao W, Jiang Y, Li Y, Wang Z, Li H, et al. Development of interactive biological web applications with R/Shiny. Brief Bioinform [Internet]. 2022 Jan 1 [cited 2024 Oct 21];23(1):bbab415. Available from: 10.1093/bib/bbab415

41. Swaminathan D, Dickinson GD, Demuro A, Parker I. Noise analysis of cytosolic calcium image data. Cell Calcium. 2020 Mar;86:102152.

42. Jacoby WG. Loess:: a nonparametric, graphical tool for depicting relationships between variables. Elect Stud [Internet]. 2000 Dec 1 [cited 2024 Oct 21];19(4):577–613. Available from: https://www.sciencedirect.com/science/article/pii/S0261379499000281

43. Cleveland WS, Devlin SJ. Locally Weighted Regression: An Approach to Regression Analysis by Local Fitting. J Am Stat Assoc [Internet]. 1988 Sep 1 [cited 2024 Oct 21];83(403):596–610. Available from: https://www.tandfonline.com/doi/abs/10.1080/01621459.1988.10478639

44. Routledge & CRC Press [Internet]. [cited 2024 Oct 21]. Statistical Models in S. Available from: https://www.routledge.com/Statistical-Models-in-S/Chambers-Hastie/p/book/9780412830402

45. Berthelot B, Grivel E, Legrand P. New Variants of DFA Based on Loess and Lowess Methods: Generalization of the Detrending Moving Average. In: ICASSP 2021 - 2021 IEEE International Conference on Acoustics, Speech and Signal Processing (ICASSP) [Internet]. 2021 [cited 2024 Oct 21]. p. 5140–4. Available from: https://ieeexplore.ieee.org/document/9414216

46. Wood SN. Generalized Additive Models: An Introduction with R, Second Edition. 2nd ed. New York: Chapman and Hall/CRC; 2017. 496 p.

47. Yang H, Stebbeds W, Francis J, Pointon A, Obrezanova O, Beattie KA, et al. Deriving waveform parameters from calcium transients in human iPSC-derived cardiomyocytes to predict cardiac activity with machine learning. Stem Cell Rep. 2022 Mar 8;17(3):556–68.

48. Mackay L, Mikolajewicz N, Komarova SV, Khadra A. Systematic Characterization of Dynamic Parameters of Intracellular Calcium Signals. Front Physiol [Internet]. 2016 Nov 10 [cited 2024 Oct 21];7. Available from: https://www.frontiersin.org/journals/physiology/articles/10.3389/fphys.2016.00525/full

49. Bootman MD, Rietdorf K, Collins T, Walker S, Sanderson M. Ca2+-sensitive fluorescent dyes and intracellular Ca2+ imaging. Cold Spring Harb Protoc. 2013 Feb 1;2013(2):83–99.

50. Knyrim M, Rabe S, Grossmann C, Gekle M, Schreier B. Influence of miR-221/222 on cardiomyocyte calcium handling and function. Cell Biosci [Internet]. 2021 Aug 17 [cited 2024 Oct 21];11(1):160. Available from: 10.1186/s13578-021-00676-4

51. Ríos E, Shirokova N, Kirsch WG, Pizarro G, Stern MD, Cheng H, et al. A preferred amplitude of calcium sparks in skeletal muscle. Biophys J [Internet]. 2001 Jan [cited 2024 Oct 21];80(1):169. Available from: https://pmc.ncbi.nlm.nih.gov/articles/PMC1301224/

52. Burridge PW, Diecke S, Matsa E, Sharma A, Wu H, Wu JC. Modeling Cardiovascular Diseases with Patient-Specific Human Pluripotent Stem Cell-Derived Cardiomyocytes. Methods Mol Biol Clifton NJ. 2016;1353:119–30.

53. Brancaccio M, Maywood ES, Chesham JE, Loudon ASI, Hastings MH. A Gq-Ca2+ axis controls circuit-level encoding of circadian time in the suprachiasmatic nucleus. Neuron. 2013 May 22;78(4):714–28.

54. Zhu MH, Jang J, Milosevic MM, Antic SD. Population imaging discrepancies between a genetically-encoded calcium indicator (GECI) versus a genetically-encoded voltage indicator (GEVI). Sci Rep [Internet]. 2021 Mar 5 [cited 2024 Oct 21];11(1):5295. Available from: https://www.nature.com/articles/s41598-021-84651-6

55. Gu X, Olson EC, Spitzer NC. Spontaneous neuronal calcium spikes and waves during early differentiation. J Neurosci Off J Soc Neurosci. 1994 Nov;14(11 Pt 1):6325–35.

56. Smith IF, Wiltgen SM, Parker I. Localization of puff sites adjacent to the plasma membrane: functional and spatial characterization of Ca2+ signaling in SH-SY5Y cells utilizing membrane-permeant caged IP3. Cell Calcium. 2009 Jan;45(1):65–76.

57. Heaney RP, Dowell MS, Hale CA, Bendich A. Calcium absorption varies within the reference range for serum 25-hydroxyvitamin D. J Am Coll Nutr. 2003 Apr;22(2):142–6.

58. Windhorst U, Johansson H, editors. Modern Techniques in Neuroscience Research [Internet]. Berlin, Heidelberg: Springer; 1999 [cited 2024 Oct 21]. Available from: https://link.springer.com/10.1007/978-3-642-58552-4

59. Sorensen J, Wiklendt L, Hibberd T, Costa M, Spencer NJ. Techniques to identify and temporally correlate calcium transients between multiple regions of interest in vertebrate neural circuits. J Neurophysiol. 2017 Mar 1;117(3):885–902.

60. Gerstein GL, Perkel DH. Simultaneously Recorded Trains of Action Potentials: Analysis and Functional Interpretation. Science [Internet]. 1969 May 16 [cited 2024 Oct 21];164(3881):828–30. Available from: https://www.science.org/doi/10.1126/science.164.3881.828

61. Smedler E, Uhlén P. Frequency decoding of calcium oscillations. Biochim Biophys Acta. 2014 Mar;1840(3):964–9.

62. Dickinson GD, Parker I. Temperature Dependence of IP3-Mediated Local and Global Ca2+ Signals. Biophys J [Internet]. 2013 Jan 22 [cited 2024 Oct 21];104(2):386. Available from: https://pmc.ncbi.nlm.nih.gov/articles/PMC3552255/

63. Gómez-Viquez NL, Guerrero-Serna G, Arvizu F, García U, Guerrero-Hernández A. Inhibition of SERCA pumps induces desynchronized RyR activation in overloaded internal Ca2+ stores in smooth muscle cells. Am J Physiol Cell Physiol. 2010 May;298(5):C1038–1046.

64. 64. findpeaks function - RDocumentation [Internet]. [cited 2024 Oct 21]. Available from: https://www.rdocumentation.org/packages/pracma/versions/1.9.9/topics/findpeaks

65. Chen TW, Wardill TJ, Sun Y, Pulver SR, Renninger SL, Baohan A, et al. Ultrasensitive fluorescent proteins for imaging neuronal activity. Nature [Internet]. 2013 Jul [cited 2024 Oct 21];499(7458):295–300. Available from: https://www.nature.com/articles/nature12354

66. Halpern ME, Rhee J, Goll MG, Akitake CM, Parsons M, Leach SD. Gal4/UAS transgenic tools and their application to zebrafish. Zebrafish. 2008;5(2):97–110.

67. E3 medium (for zebrafish embryos). Cold Spring Harb Protoc [Internet]. 2011 Oct 1 [cited 2024 Oct 21];2011(10):pdb.rec66449. Available from: http://cshprotocols.cshlp.org/content/2011/10/pdb.rec66449

68. Tan S, Zhu X, Zartman JJ, Deng Q. Low-cost Polyethylene Terephthalate Lamination Microfluidics Designs for Multiplexed Zebrafish Imaging. J Vis Exp JoVE [Internet]. 2024 Sep 27 [cited 2025 Feb 19];(211):e67313. Available from: https://app.jove.com/t/67313/low-cost-polyethylene-terephthalate-lamination-microfluidics-designs

69. Keinath NF, Waadt R, Brugman R, Schroeder JI, Grossmann G, Schumacher K, et al. Live Cell Imaging with R-GECO1 Sheds Light on flg22- and Chitin-Induced Transient [Ca(2+)]cyt Patterns in Arabidopsis. Mol Plant. 2015 Aug;8(8):1188–200.

70. Levis M, Kumar N, Apakian E, Moreno C, Hernandez U, Olivares A, et al. Microfluidics on the fly: Inexpensive rapid fabrication of thermally laminated microfluidic devices for live imaging and multimodal perturbations of multicellular systems. Biomicrofluidics. 2019 Mar;13(2):024111.

71. Dye NA, Popović M, Spannl S, Etournay R, Kainmüller D, Ghosh S, et al. Cell dynamics underlying oriented growth of the Drosophila wing imaginal disc. Dev Camb Engl. 2017 Dec 1;144(23):4406–21.

72. Giorgi FM, Ceraolo C, Mercatelli D. The R Language: An Engine for Bioinformatics and Data Science. Life Basel Switz. 2022 Apr 27;12(5):648.

73. integrate function - RDocumentation [Internet]. [cited 2024 Oct 21]. Available from: https://www.rdocumentation.org/packages/stats/versions/3.6.2/topics/integrate

74. ggplot function - RDocumentation [Internet]. [cited 2024 Oct 21]. Available from: https://www.rdocumentation.org/packages/ggplot2/versions/3.4.4/topics/ggplot

75. loess function - RDocumentation [Internet]. [cited 2024 Oct 21]. Available from: https://www.rdocumentation.org/packages/stats/versions/3.6.2/topics/loess

76. Kume S, Muto A, Inoue T, Suga K, Okano H, Mikoshiba K. Role of inositol 1,4,5-trisphosphate receptor in ventral signaling in Xenopus embryos. Science. 1997 Dec 12;278(5345):1940–3.

77. Orrenius S, Zhivotovsky B, Nicotera P. Regulation of cell death: the calcium–apoptosis link. Nat Rev Mol Cell Biol [Internet]. 2003 Jul [cited 2024 Oct 21];4(7):552–65. Available from: https://www.nature.com/articles/nrm1150

78. Härdle W, Schimek MG, editors. Statistical Theory and Computational Aspects of Smoothing [Internet]. Heidelberg: Physica-Verlag HD; 1996 [cited 2024 Oct 21]. (Müller WA, Schuster P, editors. Contributions to Statistics). Available from: http://link.springer.com/10.1007/978-3-642-48425-4

79. Filtering and Smoothing Data - MATLAB & Simulink [Internet]. [cited 2024 Oct 21]. Available from: https://www.mathworks.com/help/curvefit/smoothing-data.html

80. Rocha-Clavijo D, Suarez-Gomez D, Miranda G, Rosas NP, Alvear AEL, Braña SC, et al. Quantifying Calcium Dynamics in T Cell Populations: An Automated Analysis Framework for Antigen Fluorescence Applying Functional Anova [Internet]. Research Square; 2024 [cited 2024 Nov 20]. Available from: https://www.researchsquare.com/article/rs-5343285/v1

