## Supplemental Information for "CalciumInsights: An Open-Source, Tissue-Agnostic Graphical Interface for High-Quality Analysis of Calcium Signals"

[a]Graduate Program in Bioengineering, University of Puerto Rico at Mayagüez, Mayagüez, PR, USA
[b#]Janelia Research Campus, Howard Hughes Medical Institute, Ashburn, VA, USA
[c]Departamento de Ingeniería Química y Bioprocesos, Escuela de Ingeniería, Pontificia Universidad Católica de Chile, Santiago, Chile
[d]Department of Mathematics, University of Puerto Rico at Mayagüez, Mayagüez, PR, USA
[e]Department of Botany and Plant Pathology, Purdue University, West Lafayette, IN, USA
[f]Department of Chemical and Biomolecular Engineering, University of Notre Dame, South Bend, IN, USA
[g]Department of Biological Sciences, Purdue University, West Lafayette, IN, USA
[h]School of Mechanical Engineering, Purdue University, West Lafayette, IN, USA.
[i]College of Engineering, Texas A&M University, TX, USA
[j]Department of Biology, University of Puerto Rico at Mayagüez, Mayagüez, PR, USA
[k]Weldon School of Biomedical Engineering, Purdue University, West Lafayette, IN, USA
[l✉]Industrial Engineering Department, University of Puerto Rico at Mayagüez, Mayagüez, PR, USA
[*] Equal first-author level contribution
[#] Majority of the work was performed at the School of Mechanical Engineering and the Weldon School of Biomedical Engineering, Purdue University, West Lafayette, IN, USA.

### Supporting information

#### *Supplementary Figures*

Figure S1 represents an in-depth workflow of CalciumInsights, representing the different information flow functionalities of the software. Figures S2, S3, and S4 show the calcium traces and the metrics for the biological systems that are in Figures 6, 7, and 8, respectively, but in the y-axis log scale. In Figure S5, calcium traces with raw data and calcium traces with data smoothed using the loess method are plotted, and boxplots compare the calculated metrics with the raw and smoothed data. Table S1 presents the t-test and Wilcoxon statistical tests to compare whether there is statistical evidence that the metrics differ when calculated with the smoothed data compared to the raw data.

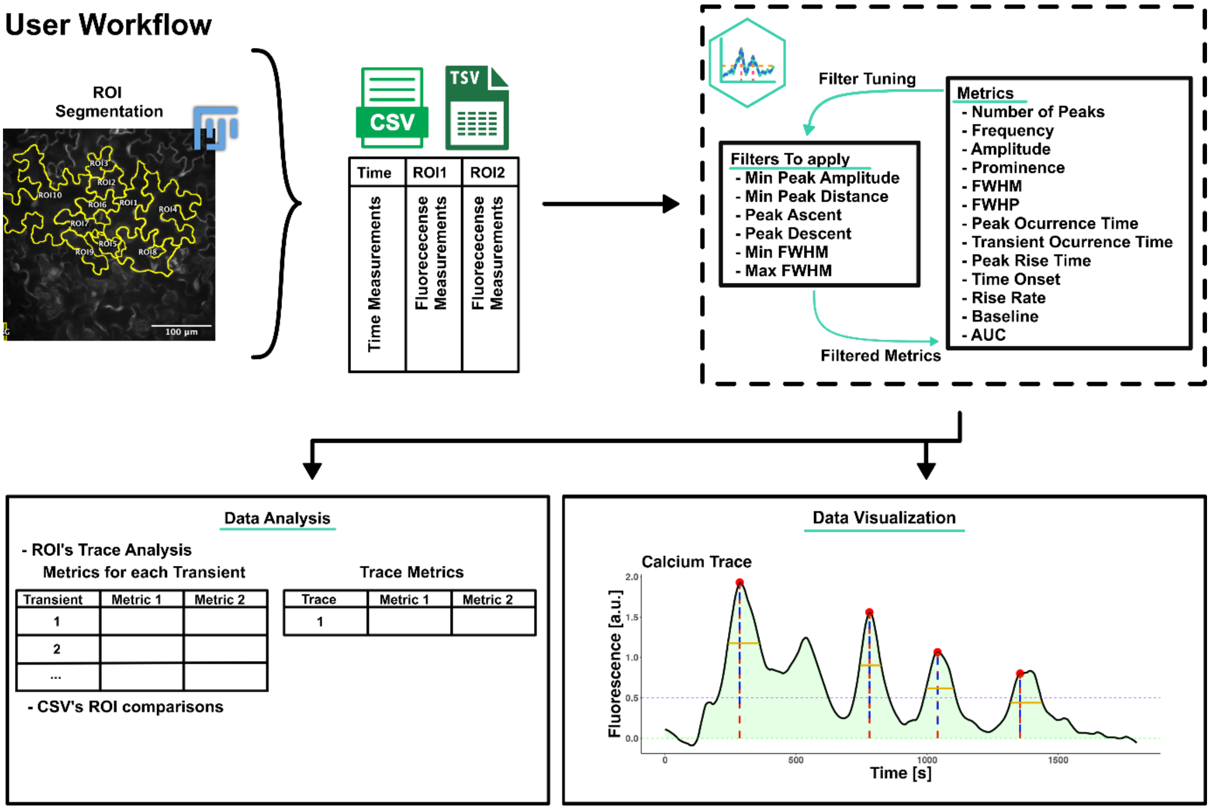


**Fig S1. In-depth workflow of CalciumInsights**

The app's flow starts with manual or automatic tissue segmentation, which exhibits $Ca^{2+}$ dynamics. The dynamics are then extracted as .csv or .tsv, which capture the dynamics as a time series, with the rows as the time and columns for each ROI. CalciumInsights can read these files, and the user can adjust the filters to provide a sense of the metrics generated by the app. Finally, the product can develop data analysis and visualization of the $Ca^{2+}$ signals extracted from a biological system.

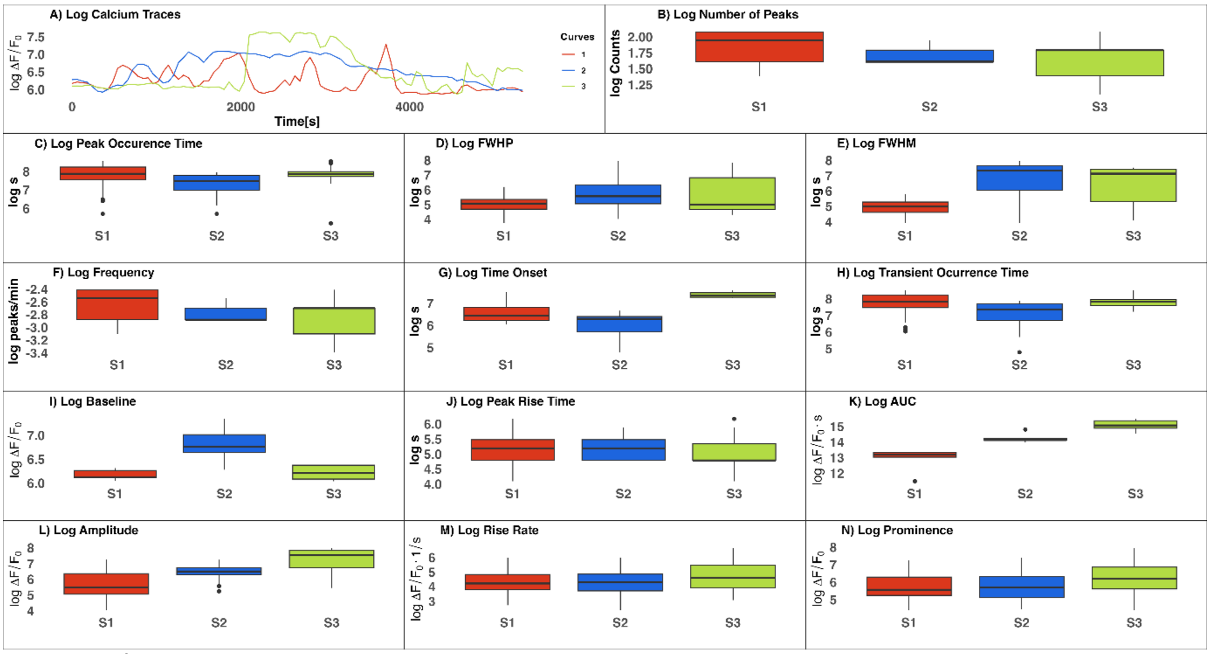


**Fig S2.** $Ca^{2+}$ traces and metrics of the epithelial layer of the resting zebrafish embryonic tail fin, 3 days post-fertilization, number of ROIs per trial = 5, and 3 trials, y-axis log scale. A) $Ca^{2+}$ Traces (one per trial to show the shape), B) Number of Peaks, C) Peak Occurrence Time, D) FWHP, E) FWHM, F) Frequency, G) Time Onset, H) Transient Occurrence Time, I) Baseline, J) Peak Rise Time, K) AUC, L) Amplitude, M) Rise Rate, and N) Prominence. Smoothness Control: 0.05, Peak Height (min): 500, FWHP (min): 40, and Prominence (min): 70.

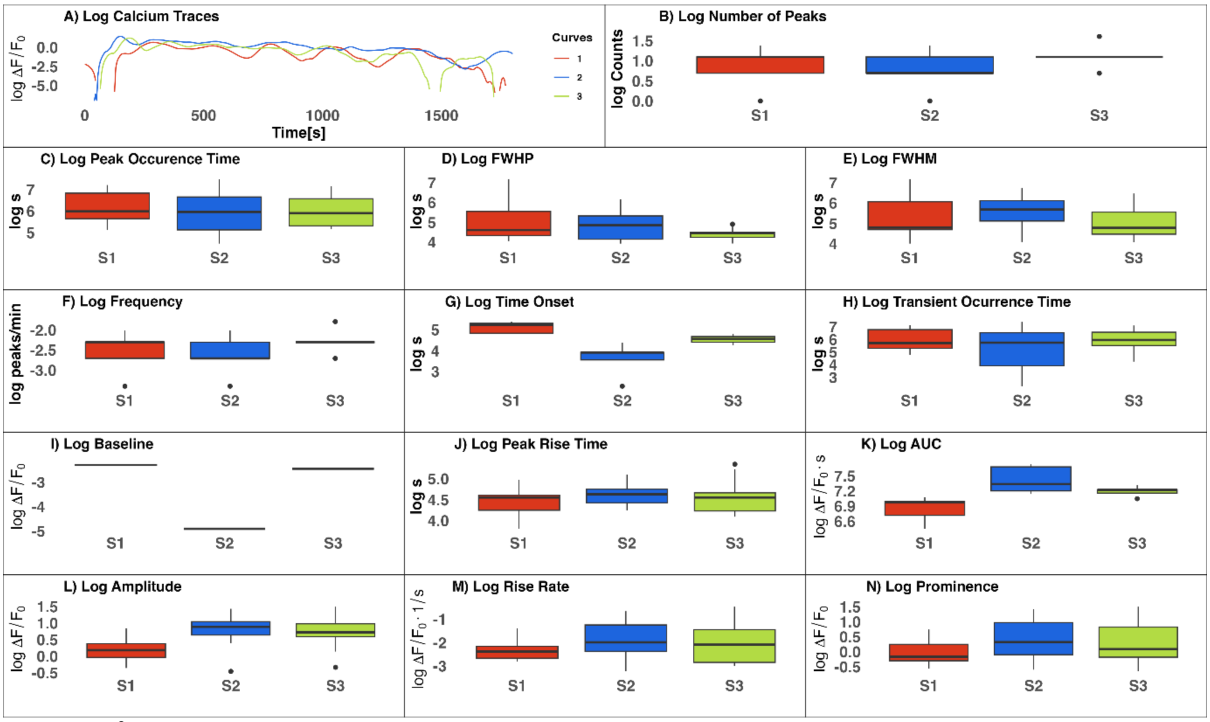


**Fig S3.** $Ca^{2+}$ transients and metrics of the cotyledon epidermal cells of *Arabidopsis thaliana*, number of ROIs per trial = 5, and 3 trials, y-axis log scale. A) Calcium Traces (one per trial to show the shape), B) Number of Peaks, C) Peak Occurrence Time, D) FWHP, E) FWHM, F) Frequency, G) Time Onset, H) Transient Occurrence Time, I) Baseline, J) Peak Rise Time, K) AUC, L) Amplitude, M) Rise Rate, and N) Prominence. Smoothness Control: 0.05, Peak Height (min): 0.5, FWHP (min): 40, and Prominence (min): 0.5.

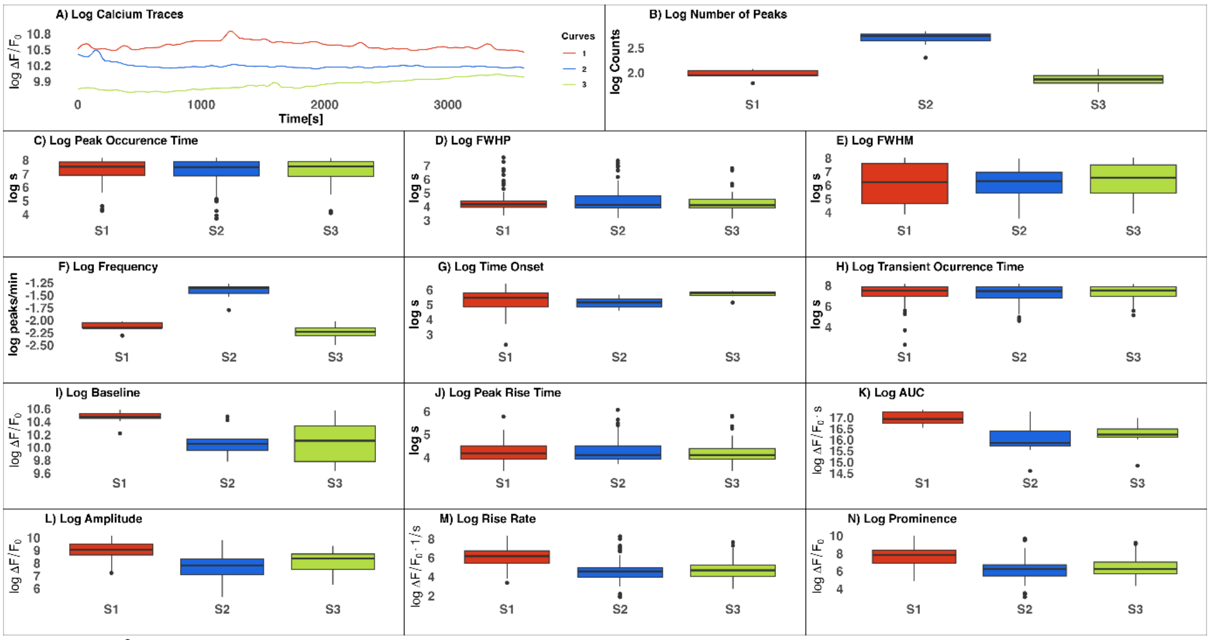


**Fig S4.** $Ca^{2+}$ transients of ex vivo wing imaginal discs from a third instar larva of *Drosophila Melanogaster,* number of ROIs per trial = 10, and 3 trials, y-axis log scale. A) Calcium Traces (one per trial to show the shape), B) Number of Peaks, C) Peak Occurrence Time, D) FWHP, E) FWHM, F) Frequency, G) Time Onset, H) Transient Occurrence Time, I) Baseline, J) Peak Rise Time, K) AUC, L) Amplitude, M) Rise Rate, and N) Prominence. Smoothness Control: 0.04, Peak Height (min): 17000, Peak Ascend: 3, Peak Descend: 3, Min Peak Distance: 36.

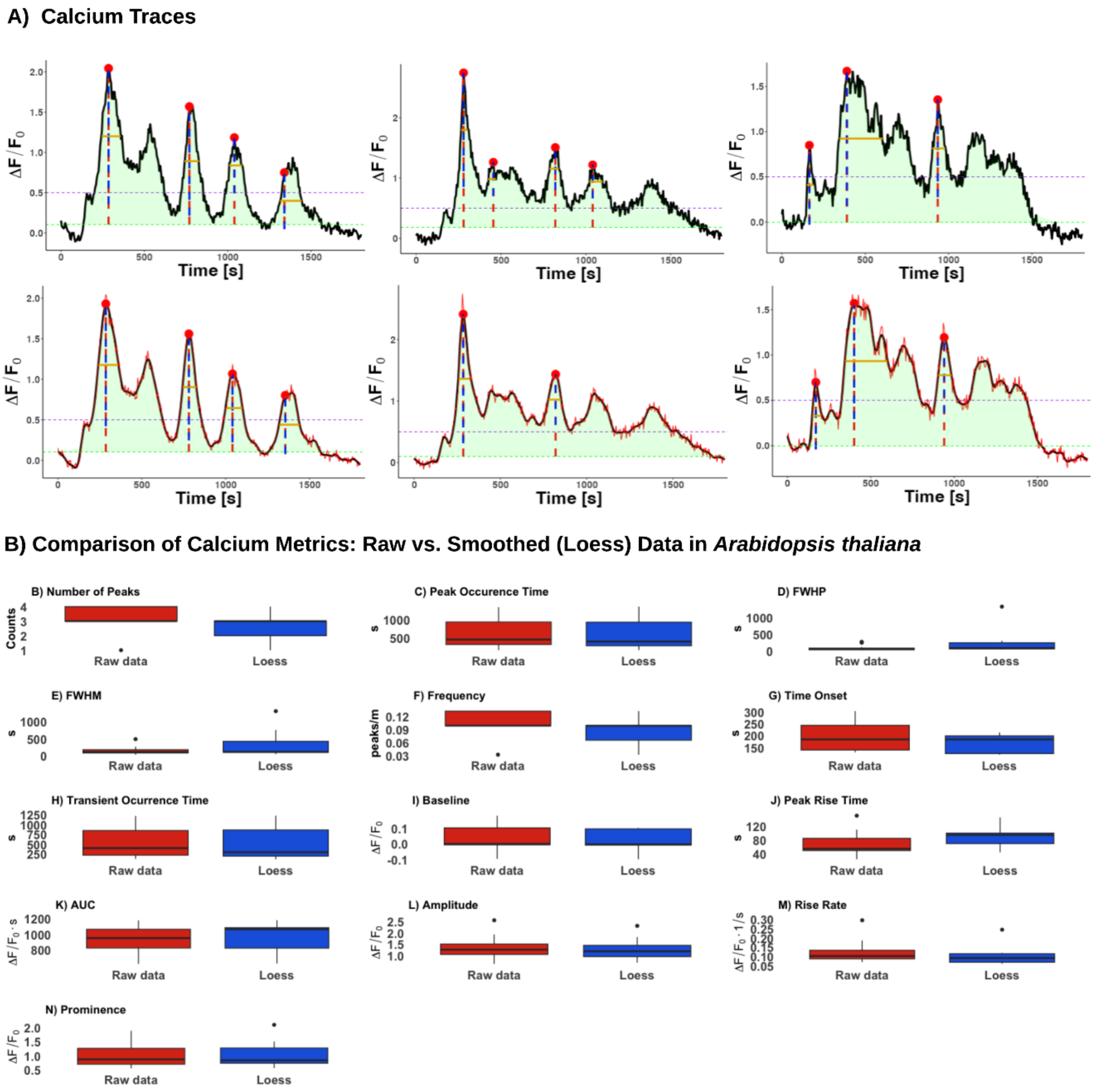


**Fig S5.** A) A biological replicate of *Arabidopsis thaliana* was taken, and calcium metrics were calculated from raw and smoothed data using the Loess method. B) Boxplots were generated to compare the visual and descriptive differences between the calcium metrics calculated from the raw data and those smoothed with Loess.

**Results of hypothesis testing for comparing metrics calculated from raw and smoothed data.**

| Metric | t statistic | T_Test_P_Value | W | Wilcoxon_Test_P_Value |
|---|---|---|---|---|
| **Amplitude** | 0.302 | 0.764 | 104.0 | 0.782 |
| **Peak_Occurence_Time** | 0.103 | 0.918 | 98.5 | 0.981 |
| **Prominence** | -0.193 | 0.848 | 88.0 | 0.678 |
| **FWHP** | -1.358 | 0.197 | 58.0 | 0.072 |
| **FWHM** | -1.536 | 0.146 | 73.0 | 0.268 |
| **Peak_Rise_Time** | -1.890 | 0.069 | 55.0 | 0.051 |
| **Transient_Occurrence_Time** | 0.264 | 0.793 | 108.5 | 0.628 |
| **Rise_Rate** | 1.042 | 0.306 | 126.0 | 0.197 |
| **Time_Onset** | 0.834 | 0.433 | 16.5 | 0.463 |
| **Frequency** | 0.534 | 0.607 | 15.5 | 0.583 |
| **Baseline** | 0.301 | 0.771 | 16.0 | 0.530 |
| **Number_of_Peaks** | 0.534 | 0.607 | 15.5 | 0.583 |
| **AUC** | -0.198 | 0.847 | 10.0 | 0.676 |

**Table S1.** The results of the hypothesis tests conducted for each calculated metric, aimed at assessing whether there are significant differences between the metrics obtained from the raw and smoothed data. From it we can see that none of them are statistically significant.

#### *Metadata*

| **Species** | ***Arabidopsis*** | ***Danio rerio*** | ***Drosophila melanogaster*** |
|---|---|---|---|
| Sample description | Arabidopsis 7-d old cotyledon epidermal cells treated with 1 uM flg22 | Tg(krt:gal4, UAS:GCaMP6f) live zebrafish embryos, 3 days post fertilization | *Drosophila* wing disc dissected out of 3rd instar wandering larva 6 days after egg laying |
| Sample preparation | Individual cotyledons were excised from seedlings and mounted in custom-made chambers. The abaxial side was imaged with an inverted microscope. | Tricaine (MS-222) added to medium with live embryos to final concentration of 164 mg/L. At 60min, latrunculin A was added to final concentration of 2uM | The dissected-out disc was treated with 1 mM Yoda1 for 1 hour to activate the Piezo channels |
| Mounting medium | water | E3 (using PETL laminate microfluidics device) | Grace's insect media with 20 ecdysones |
| Image width in pixels (X) | 512 | 512 | 512 |
| Image height in pixels (Y) | 512 | 512 | 512 |
| Number of slices (Z) | 1 | 27 pre-treatment, 24 post-treatment (compressed into one slice via max intensity projection) | 1 |
| Number of channels (C) | 1 | 2 (only one analyzed) | 1 |
| Number of frames (T) | 360 | 54 @ 65s per frame (pre-addition) + 90 @ 60s per frame (post-addition) | 301 |
| Pixel size XY (micron) | 0.667 | 1.24 | 0.34 |
| Voxel size Z (micron) | NA | 2.33 (compressed via max intensity projection) | NA |
| Time interval (second) | 5 | 60 | 10 |
| Microscope | Olympus/Andor spinning disk confocal | Nikon A1Rmp | Nikon Eclipse Ti confocal microscope with a Yokogawa spinning disc |
| Acquisition software | MetaMorph | NIS Elements | MetaMorph |
| Acquisition objective | Olympus 20X 0.4–NA Plan C Achromat | Plan Apo VC 20x/0.75 DIC N2 | Nikon, Plan Fluor, 40x 1.30-NA Oil, DIC H/N2 |

| | | | |
|---|---|---|---|
| Channel description (e.g. fluorophore, labeled protein or cell type) | R-GECO1 | 488nm/FITC/Green, 560/TRITC/Red (unused) | GCaMP6f |
| Light source (e.g. laser 488nm) | laser 561nm | Laser 488nm, 560nm | 488 nm laser |
| Emission filter | 610/37-nm | 488/525 +/- 50 (green), 560/595 +/- 50 (red, unused) | 525/50 nm |

#### *Linear Regression*

Linear regression is a statistical method used to model the relationship between a dependent variable and one or more independent variables by fitting a linear equation to the collected data. The general formulation of linear regression for this model is:

$$\hat{y} = b_0 + b_1 x, \qquad (1)$$

where $\hat{y}$ represents the predicted fluorescent value and $b_0$ is the intercept or constant term of the model, which represents the value of $\hat{y}$ when $x$ is equal to zero. $x$ denotes the raw fluorescence data, serving as the independent variable, $b_0$ and $b_1$ are the weights adjusted to minimize the difference between observed and predicted values (1).

Linear regression is employed to analyze the behavior of raw and smoothed fluorescence data, enabling the assessment of how well the smoothed curve from the Loess method fits the original data. The fit is estimated to be optimal when the smoothed fluorescence values approach the linear regression model more closely (2).

$R^2$ is a coefficient indicating the proportion of variability in the dependent variable that can be explained by the independent variable in a linear regression model. This value ranges between 0 and 1, where 0 signifies that the independent variable does not explain any variability in the dependent variable, while 1 indicates a perfect fit of the model (3).

The formula for the coefficient of determination$R^2$ is given by:

$$Error\ sum\ of\ squares = \sum_{i=1}^{n}(y_i - \hat{y}_i)^2, \qquad (2)$$

$$Total\ sum\ of\ squares = \sum_{i=1}^{n}(y_i - \bar{y}_i)^2, \qquad (3)$$

$$R^2 = 1 - \frac{Error\ sum\ of\ squares}{Total\ sum\ of\ squares} \qquad (4)$$

where, in this case, $y_i$ is the raw fluorescence [a.u.], the $\hat{y}_i$ is the smoothed fluorescence [a.u.] generated by the Loess algorithm and $\bar{y}$ is the mean of the raw fluorescence [a.u.].

**Savitzky-Golay**: The Savitzky-Golay method is a data smoothing technique used in digital signal processing. It involves fitting low-degree polynomials to successive subsets of adjacent points using the least squares method. Its main advantage is that it preserves key features of the original signal, such as maxima, minima, and peak width, making it particularly useful for spectroscopic analysis and time series processing. It was proposed by Abraham Savitzky and Marcel J. E. Golay in 1964 (4).

#### References


1. 2.3 - The Simple Linear Regression Model | STAT 462 [Internet]. [cited 2024 Oct 21]. Available from: https://online.stat.psu.edu/stat462/node/93/
2. What Is Linear Regression? | IBM [Internet]. 2021 [cited 2024 Oct 21]. Available from: https://www.ibm.com/topics/linear-regression
3. 2.5 - The Coefficient of Determination, r-squared | STAT 462 [Internet]. [cited 2024 Oct 21]. Available from: https://online.stat.psu.edu/stat462/node/95/
4. Konstantinovsky T. Medium. 2024 [cited 2025 Mar 24]. Introduction to the Savitzky-Golay Filter: A Comprehensive Guide (Using Python). Available from: https://medium.com/pythoneers/introduction-to-the-savitzky-golay-filter-a-comprehensive-guide-using-python-b2dd07a8e2ce
